# Lipopolysaccharide truncation and restoration drives a trade-off in resistance to two phages in *Pseudomonas aeruginosa*

**DOI:** 10.64898/2026.01.20.700494

**Authors:** Ellie J Tong, Luis M Bolaños, Julie Fletcher, Robyn Manley, Christian Fitch, Christina Bugert, Jacob Sturgess, James W Sheffield, Rosie Allen, James Graham, Natalie Whitehead, Tom Ireland, Steven L Porter, Ben Temperton

**Affiliations:** Biosciences, Faculty of Health and Life Sciences, University of Exeter, Stocker Road, Exeter EX4 4QD, UK; Institute of Quantitative Biology, Biochemistry and Biotechnology, University of Edinburgh, Alexander Crum Brown Road, The King’s Buildings, Edinburgh EH9 3FF; University of Sheffield, Western Bank, Sheffield, S10 2TN; Churston Ferrers Grammar School, Greenway Road, Brixham, Devon, TQ5 0LN; Exeter Science Centre, Kaleider Studios, 45 Preston Street, Exeter EX1 1DF, UK

**Keywords:** *Pseudomonas aeruginosa*, phage therapy, phage receptor, phage resistance, receptor modification, lipopolysaccharide

## Abstract

Phage therapy is a promising treatment for multidrug resistant bacterial infections, and for patients no longer able to tolerate antibiotic treatments. A major challenge for phage therapy is emergent phage resistance, which target bacteria acquire by structurally modifying or masking phage receptors to prevent adsorption. Functionally diverse phage cocktails that target a broad range of receptors are less prone to resistance as there is a higher fitness cost associated with modifying multiple receptors. Expanding phage libraries with well-characterised phages that target a broad range of receptors would aid in timely and strategic design of functionally diverse phage cocktails. Here, we aimed to isolate phages targeting novel receptors by enriching wastewater samples on a *Pseudomonas aeruginosa* PAO1 Δ*pilA* Δ*galU* unmarked deletion mutant lacking O-antigen, outer core lipopolysaccharide (LPS) and type IV pili (T4P) - the three most common *Pseudomonas* phage receptors. This led to the isolation of a novel phage, named *Vale*. Vale was predicted to bind the LPS inner core as it could only infect strains with truncated LPS, suggesting that the outer core LPS blocks Vale from accessing the inner core. We identified a trade-oM in resistance to Vale and another phage, *Tor,* that targets the LPS outer core, mediated by host-derived LPS modifications. The PAO1 host evolved resistance to Tor by 100-200kb genomic deletions, which resulted in LPS truncation and sensitivity to Vale. Complete LPS restoration in the Δ*pilA* Δ*galU* mutant conferred resistance to Vale and sensitivity to Tor in two out of three replicates. Combined treatment with Tor and Vale delayed the emergence of resistance in PAO1 for at least three times longer than individual phage treatments. This study provides an example of how using phage receptors to strategically design phage cocktails can minimise the likelihood of emergent phage resistance.

**Graphical abstract:** A summary of the LPS modifications, genomic mutations and phage susceptibilities of Tor and Vale resistant mutants. “Parent strain” refers to PAO1 Δ*hsdR.* Δ*pilA* Δ*galU* refers to PAO1 Δ*hsdR* Δ*pilA* Δ*galU.* The parent strain gains resistance to Tor via LPS truncation associated with 100-200 kb genomic deletions, resulting in sensitivity to Vale. Δ*pilA* Δ*galU* gains resistance to Vale by restoring its LPS, conferring sensitivity to Tor in two out of three repeats. As resistance to one phage sensitises bacteria to the other, combined treatment with both phages suppresses phage resistance for longer than individual treatments.

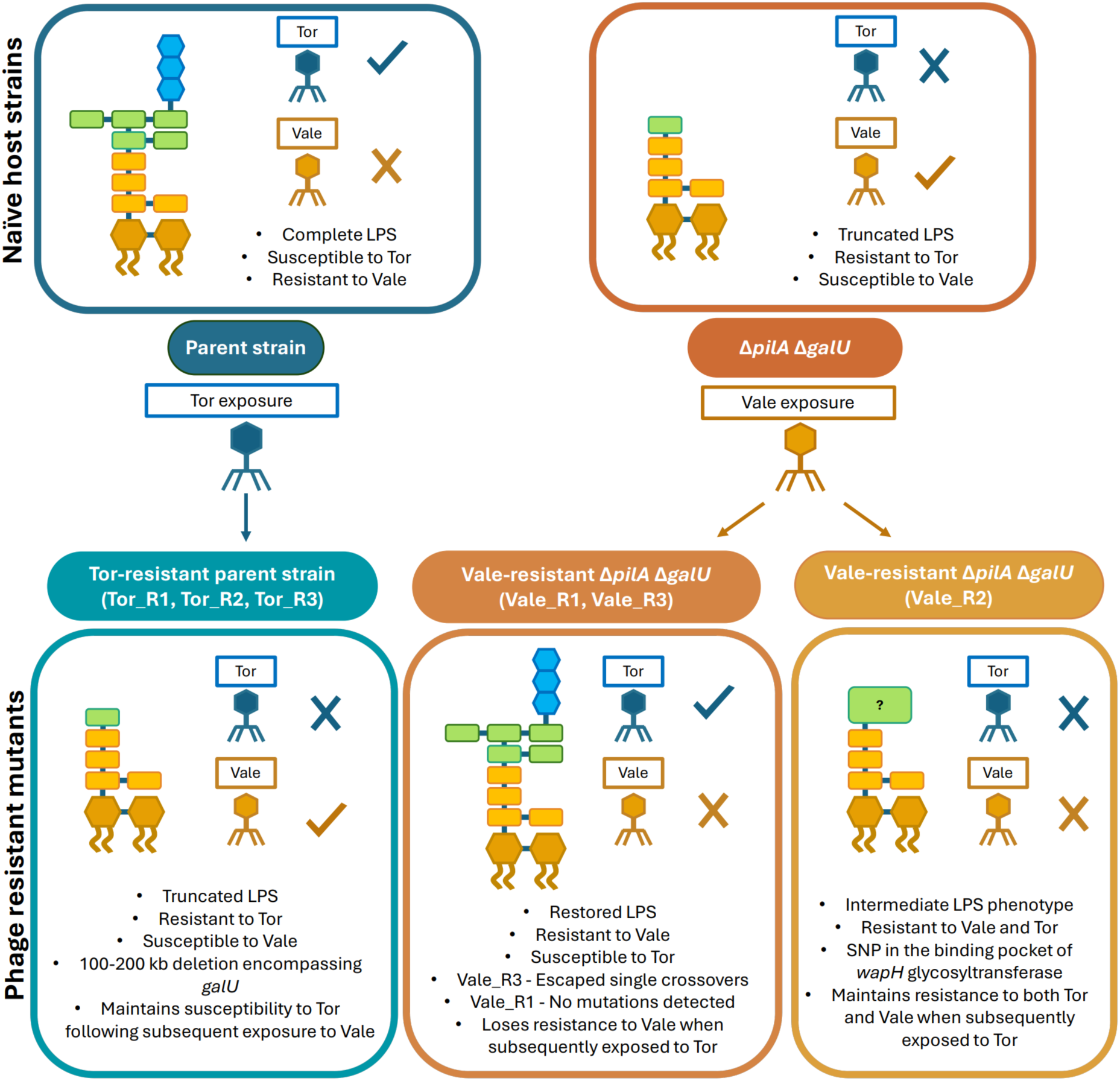

## Introduction

*Pseudomonas aeruginosa* is an opportunistic pathogen that causes severe infections in immunocompromised individuals, including cystic fibrosis (CF), bronchiectasis, burn wound, and transplant patients^1–3^. A large, plastic genome affords *P. aeruginosa* broad intrinsic and acquired antibiotic resistance, with up to half of intensive care isolates exhibiting muti-drug resistance (MDR)^4^, most frequently associated with chronic respiratory infections^5,6^. While the World Health Organisation (WHO) designates *P. aeruginosa* a critical priority for new antibiotics, regulatory and economic barriers have stalled their development^7,8^. Phage therapy is a promising alternative or adjunctive treatment, using lytic bacteriophage (phage), to clear infections^9^.

Lytic phage infection begins with adsorption to the bacterial host, mediated by an extremely specific interaction between tail fibre binding domains and bacterial surface receptors^10^. A major benefit of phage therapy over broad-range antibiotics is that high receptor specificity narrows phage host range, allowing targeted killing of pathogens without harming commensal microbes^11^. However, a narrow host range also limits the number of phages that can infect each clinical isolate, and often specific combinations of phages (cocktails) must be tailored to each patient strain^12^. Additionally, target pathogens can rapidly evolve phage resistance by modifying or masking phage receptors to prevent phage adsorption, both *in vitro*^13,14^ and *in vivo*^15^. As a result, bacteria can acquire cross-resistance to multiple phages that use the same receptor through a single receptor modification^16^; this is particularly problematic when selecting phages to treat *P. aeruginosa* infections, as there are relatively few known *Pseudomonas* phage receptors^17^. Furthermore, modifying phage receptors compromises their normal function, meaning phage resistance is frequently associated with fitness trade-offs^15,18,19^. Overall, receptor specificity is an important consideration when designing phage cocktails with sufficient ‘breadth’ to effectively target specific clinical strains and sufficient ‘depth’ to minimise the risk of phage resistance^12,20,21^.

Functionally diverse phage cocktails that target a variety of different receptors are less prone to emergent host resistance, as there is a higher fitness cost associated with modifying multiple receptors^22–25^. For instance, one study found that treatment of *Klebsiella pneumoniae* with a three-phage cocktail targeting three distinct receptors delayed phage resistance for at least 40 hours, compared to 15-30 hours for a two-phage treatment. By contrast, a functionally redundant three-phage cocktail, with two phages targeting the same receptor, triggered earlier and more frequent phage resistance^22^. Targeting different regions or structural morphologies of the same receptor can incur exploitable fitness trade-offs, as modifying the receptor to resist one phage may sensitise the bacteria to another ^23,26^. This was demonstrated when a five-phage *P. aeruginosa* cocktail, with phages targeting distinct regions of the lipopolysaccharide (LPS), suppressed emergence of phage resistance for up to five days^26^. Additionally, “phage steering” approaches select phages binding receptors involved in drug resistance or virulence to exploit the trade-offs associated with receptor modification and steer the population to a more treatable and benign state^15,18,19^. For example, resistance to OMKO1, one of the few *Pseudomonas* phages known to target the antibiotic efflux porin OprM, resulted in impaired antibiotic efflux and increased susceptibility to several antibiotics^18,27^. Thus, strategically designed phage cocktails targeting specific and diverse receptors can suppress phage resistance and capitalise on the associated fitness trade-offs^13,20^.

The most frequently reported *P. aeruginosa* phage receptors are the type IV pili (T4P) and LPS^17,28^. The LPS is characterised into four parts: lipid A, inner core, outer core and O-antigen, with phages most commonly binding the outer core and O-antigen (Figure 1). Both LPS and T4P are important virulence factors involved in motility, adhesion and biofilm formation ^29–32^, and LPS structure influences membrane permeability to various antibiotics^32^. Phages utilising these receptors have been successfully used to treat *P. aeruginosa* infections *in vivo*^28^ with anti-virulence and phage steering effects demonstrated *in vitro*^28,33^. However, phage resistance readily emerges via loss or glycosylation of T4P and LPS modifications, limiting the efficacy of phage cocktails targeting these receptors alone ^28,34–36^. Additionally, receptor identification is rarely a routine step of phage characterisation, primarily because designing personalised phage cocktails is time-sensitive and already entails numerous stages of characterisation and purification^20^. Characterising and expanding phage libraries to target a broader range of receptors would assist in the development of personalised and functionally diverse phage cocktails that are less prone to resistance.

**Figure 1:**
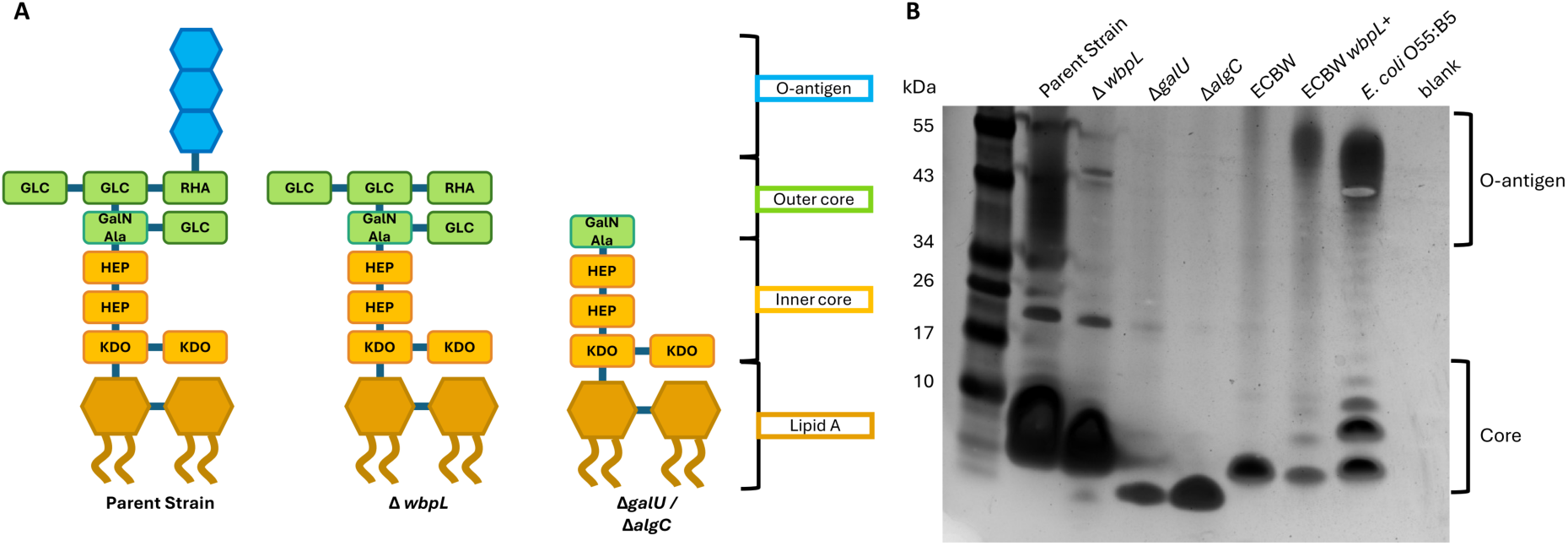
**[A]** The predicted structure of LPS deletion mutants. The parent strain possesses a complete LPS core with O-antigen. Δ*wbpL* possesses a complete core, devoid of O-antigen^39^. Δ*galU* and Δ*algC* are devoid of O-antigen and have a truncated LPS core^45,79^. Residue abbreviations are: 3-deoxy-D-manno-ocr-2-ulosonic acid (KDO), heptose (HEP), 2-(2-L-alanyl)-2 deoxy-D-galactosamine (GalNAla), glucose (GLC), rhamnose (RHA). **[B]** LPS extractions from LPS mutants run on a 4-12% bis-tris SDS-PAGE gel and silver stained. From right to left, the lanes show LPS extractions from (1) the parent strain, (2) Δ*wbpL,* (3) Δ*galU,* (4) Δ*algC,* (5) *E. coli* BW25113 and (6) *E. coli* BW25113 *wbpL*+, (7) *E. coli* O55:B5 commercial LPS (Sigma-Aldrich), and (8) nuclease-free water blank.

Here, we constructed a panel of unmarked deletion mutants lacking known *P. aeruginosa* phage receptors (Table 1) as a tool for characterising the receptors targeted by *P. aeruginosa* phages in the Citizen Phage Library (CPL)^37^. We also developed a Δ*pilA* Δ*galU* mutant, which lacks O-antigen, outer core LPS, and T4P to use as an enrichment host for isolating phages that target more unique receptors, particularly OprM, due to the high clinical value of phages capable of phage steering to restore drug susceptibility. To improve enrichment yields and facilitate phage propagation, a previously developed promiscuous PAO1 Δ*hsdR*, which lacks the defensive type I restriction endonuclease^38^, served as the parent strain for all receptor deletions (for clarity, unless otherwise stated, *ΔhsdR* is assumed throughout, i.e. Δ*pilA* Δ*galU* is PAO1 Δ*hsdR* Δ*pilA* Δ*galU* and “parent strain” refers to PAO1 *ΔhsdR*). While no phages targeting OprM were identified, a novel phage, Vale, was found to target the inner core LPS. Vale could only infect strains with truncated LPS, either via gene deletion or via emergent resistance against a phage targeting the outer core (Tor). We identified a trade-oM in resistance to Vale and Tor, where resistance to one phage sensitises the bacteria to the other via LPS modifications. Combined treatment with Vale and Tor effectively suppressed phage resistance for up to three times longer than individual phage treatments.

**Table 1:**
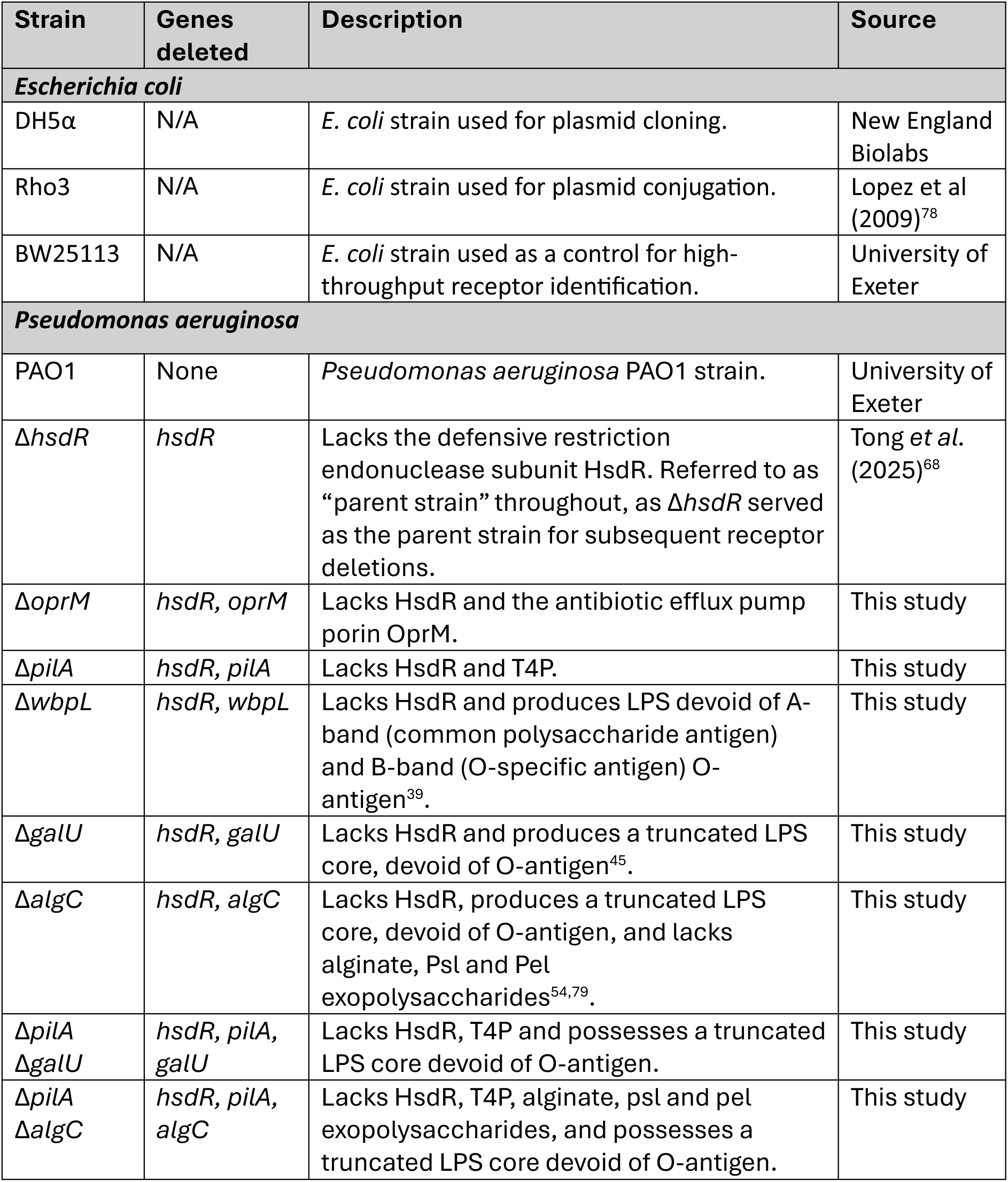
The bacterial strains used in this study.

## Results

### Development of a panel of unmarked deletion mutants lacking common phage receptors

A panel of unmarked deletion mutants lacking previously reported *Pseudomonas* phage receptors was generated by two-step allelic exchange^38^ in the promiscuous PAO1 strain Δ*hsdR* (Table 1). Successful deletions were identified by a check PCR amplifying the region surrounding the target gene (Supplementary Figure 1) and confirmed with Sanger sequencing of the PCR product (Supplementary Materials). SDS-PAGE of LPS extractions confirmed that deletions of LPS biosynthesis genes had the desired phenotypic effects (Figure 1). Most gene deletions had no significant effect on doubling time or carrying capacity compared to Δ*hsdR* (Supplementary Figure 2 and Supplementary Table 3). Δ*algC* had a 2-fold longer doubling time and Δ*wbpL* demonstrated population decline soon after reaching carrying capacity (Supplementary Table 3). Curiously, Δ*pilA* Δ*galU* possessed a 1.7-fold higher carrying capacity than Δ*hsdR*.

### Outer core LPS and T4P are common receptors of Citizen Phage Library phages

The receptors of 50 previously isolated *Pseudomonas* phages within the CPL were identified by screening their host range across the panel of receptor deletion mutants (Table 1). When a phage produced a zone of lysis on the parent strain but not on a given receptor mutant, this indicated that the deleted structure was required as a phage receptor (Figure 2). Thirty-six phages did not infect Δ*galU* or Δ*algC*, but did infect Δ*wbpL,* indicating that they require the LPS outer core as a receptor. Four phages did not infect Δ*galU,* Δ*algC* or Δ*wbpL,* suggesting that they require O-antigen. Ten phages could not infect Δ*pilA*, suggesting that they use T4P as their primary receptor. Out of the ten phages that use T4P, six could not infect Δ*wbpL*. *wpbL* is involved in the synthesis of A-and B-band O-antigens^39^, but has also been shown to glycosylate T4P^34,40^, and thus it is likely that these phages require the glycosylated form of T4P, rather than using the O-antigen as a secondary receptor. Overall, no previously isolated phage could infect the Δ*pilA* Δ*galU* and Δ*pilA* Δ*algC* mutants, meaning all 50 phages required T4P, outer core LPS, or O-antigen as their receptor.

**Figure 2:**
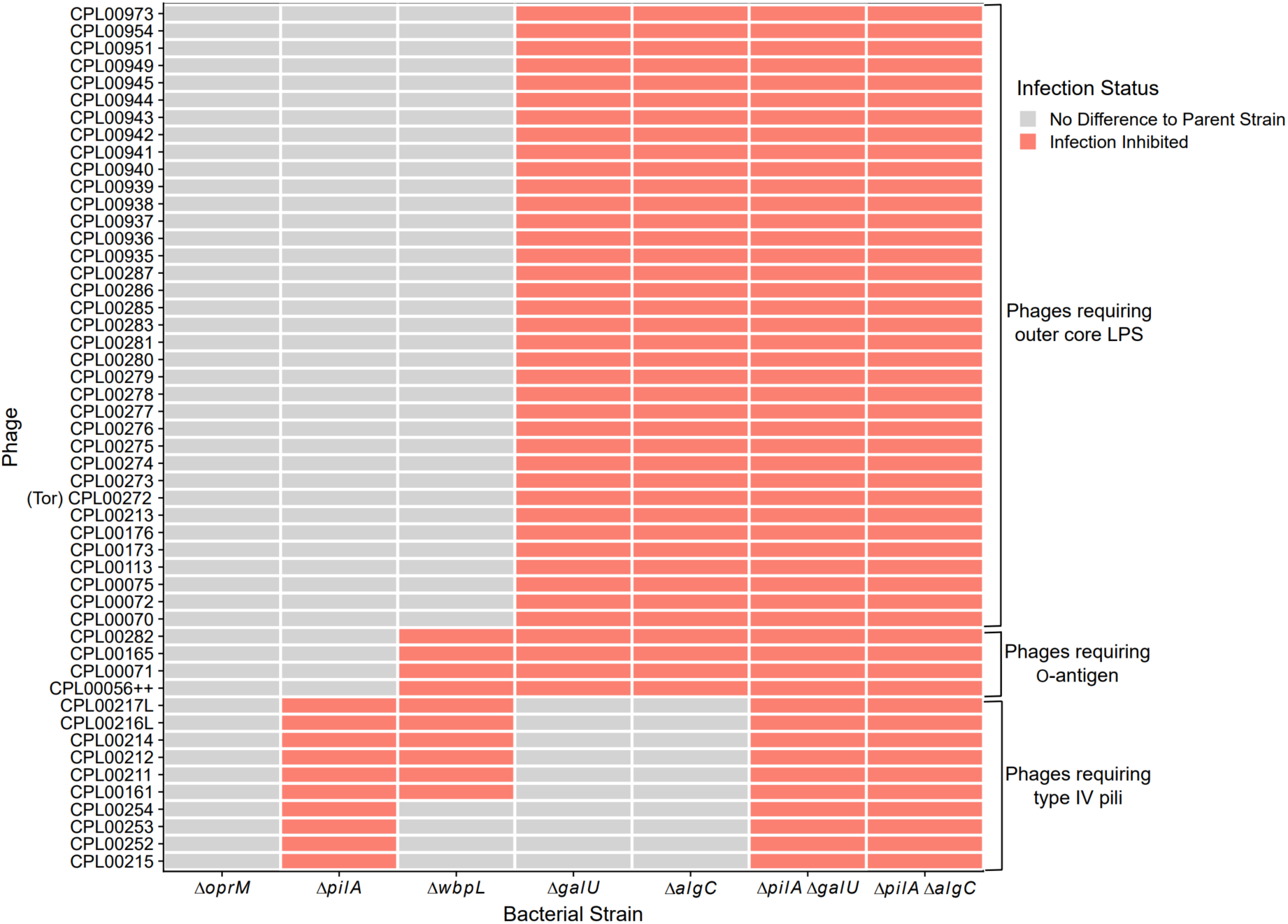
The host range of 50 Citizen Phage Library (CPL) *P. aeruginosa* phages across the panel of receptor mutants. The 50 phages were originally isolated on PAO1 or clinical *Pseudomonas* strains (Supplementary Materials). Inhibited infection (red) on a given mutant compared to the parent strain indicates that the deleted structure is a receptor required for infection.

### Using Δ*pilA* Δ*galU* as an enrichment host yields phages that bind alternative receptors

To isolate phages using novel receptors, Δ*pilA* Δ*galU* and Δ*pilA* Δ*algC* mutants, which lack the LPS outer core, O-antigen and T4P, were used as enrichment hosts. Initially, enrichment from 95 wastewater samples on Δ*pilA* Δ*galU* and Δ*pilA* Δ*algC* yielded one phage, CPL01271, on Δ*pilA* Δ*galU*. Another 95 wastewater samples were enriched on Δ*pilA* Δ*galU* in the presence of trimethoprim to encourage the upregulation of OprM expression, increasing the chance of isolating OprM-targeting phages. Two and eight phages were isolated in the presence of 25 µg/mL (MIC×10) and 2.5 µg/mL (At MIC) trimethoprim, respectively. The sequence of one of these phages, CPL01283, assembled as two distinct phages, herein referred to as CPL01283_1 and CPL1283_2. Taxonomic classification via average nucleotide similarity suggested most were *PhiKZ-like* viruses or *Pbunaviruses,* except CPL01282, CPL01283_1 and CPL01283_2, which were unknown species with < 80% identity to any phage from the NCBI genbank or CPL databases (Table 2, Supplementary Figure 3).

**Table 2:** The phages isolated on *ΔpilA ΔgalU.* The closest relatives were identified from the NCBI Genbank database and intergenomic distance calculated in PhageClouds^69^.

| Phage | Isolation host and conditions | Predicted receptor | Species | Closest relative |  |  |
| --- | --- | --- | --- | --- | --- | --- |
|  |  |  |  | Phage name | Accession number | Distance |
| CPL01271 | $\Delta pilA \Delta galU$ | Unknown | *Pbunavirus* | LBL3 | NC_011165 | 0.0321 |
| CPL01275 | $\Delta pilA \Delta galU$ 25 µg/mL trimethoprim | Unknown | *PhiKZvirus* | KTN4 | KU521356 | 0.0121 |
| CPL01276 (Vale) | $\Delta pilA \Delta galU$ 25 µg/mL trimethoprim | Inner core LPS | *Pbunavirus* | vB_PaeM_SMS29 | MN615702 | 0.0191 |
| CPL01277 | $\Delta pilA \Delta galU$ 2.5 µg/mL trimethoprim | Unknown | *PhiKZvirus* | KTN4 | KU521356 | 0.0108 |
| CPL01278 | $\Delta pilA \Delta galU$ 2.5 µg/mL trimethoprim | Unknown | *PhiKZvirus* | PaZh_1 (unverified organism) | MN871489 | 0.0097 |
| CPL01279S | $\Delta pilA \Delta galU$ 2.5 µg/mL trimethoprim | Exopolysaccharide | *Pbunavirus* | vB_PaeM_USP_1 | NC_050149 | 0.0368 |
| CPL01279L | $\Delta pilA \Delta galU$ 2.5 µg/mL trimethoprim | Inner core LPS | *Pbunavirus* | vB_PaeM_SMS29 | MN615702 | 0.0318 |
| CPL01280 | $\Delta pilA \Delta galU$ 2.5 µg/mL trimethoprim | Inner core LPS | *Pbunavirus* | vB_PaeM_SMS29 | MN615702 | 0.0318 |
| CPL01282 | $\Delta pilA \Delta galU$ 2.5 µg/mL trimethoprim | Inner core LPS | Unclassified | BUCT566 | NC_074664 | 0.0688 |
| CPL01283_1 | $\Delta pilA \Delta galU$ 2.5 µg/mL trimethoprim | Unknown | Unclassified | BUCT566 | NC_074664 | 0.0687 |
| CPL01283_2 | $\Delta pilA \Delta galU$ 2.5 µg/mL trimethoprim | Unknown | Unclassified | AUS531phi | MN585195 | 0.0679 |
| CPL01284 | $\Delta pilA \Delta galU$ 2.5 µg/mL trimethoprim | Unknown | *PhiKZvirus* | PA02 | AP019418 | 0.0143 |

Screening the novel phages against the deletion mutant panel indicated that none required OprM as their receptor, as all could infect the Δ*oprM* mutant (Figure 3). CPL01271 formed zones of lysis on all the receptor mutants (Figure 3) and no SNPs were identified from short-read sequences of two CPL01271-resistant mutants. Therefore, CPL01271 uses a currently unknown receptor that is not O-antigen, outer core LPS, T4P or OprM. CPL01284 did not consistently form clear zones of lysis on LPS mutants, despite originally being isolated on Δ*pilA ΔgalU.* Four phages infected strains with wildtype LPS and Δ*galU* mutants but had varied ability to infect Δ*algC* and Δ*wbpL* mutants. The remaining four phages only infected strains with truncated LPS. CPL01279L and CPL01280 only infected strains lacking O-antigen (Δ*wbpL,* Δ*galU* and Δ*algC*), and CPL01276 (Vale) and CPL01282 only infected strains that lacked the outer core LPS (Δ*galU* and Δ*algC*). This suggests that these phages bind part of the inner core LPS, which is only exposed when O-antigen is lost, or the outer core is truncated. To our knowledge only one phage, the dsRNA phage PhiYY, has been predicted to use the *P. aeruginosa* LPS inner core previously. Thus, these *Pbunaviruses* are the first dsDNA phages identified to use the *P. aeruginosa* inner core LPS.

**Figure 3:**
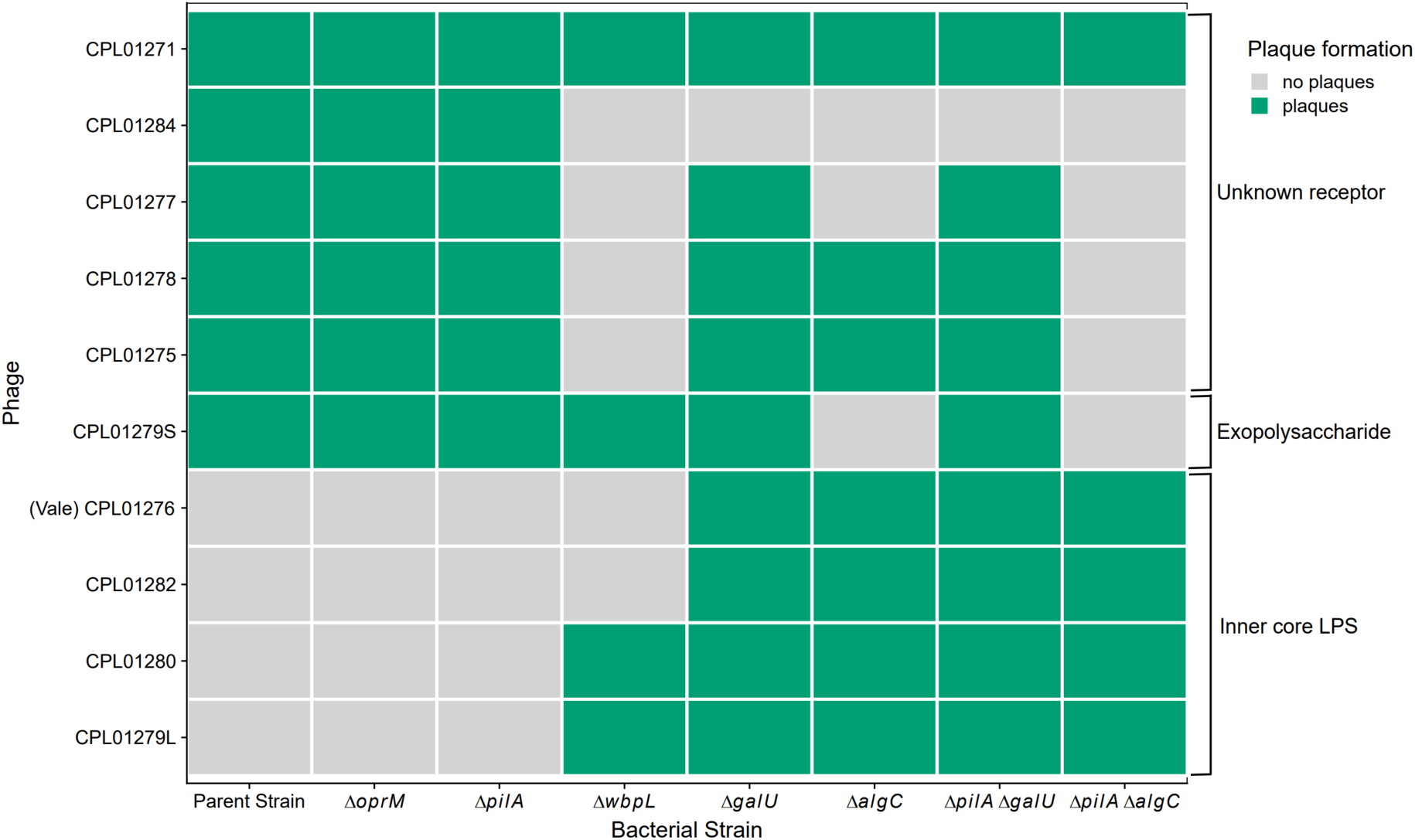
The host range of phages isolated on Δ*pilA* Δ*galU* across the panel of receptor mutants. The parent strain for all receptor mutants was the promiscuous strain PAO1 Δ*hsdR.* Green squares indicate that the phage formed a zone of lysis on the bacterial strain.

### LPS truncation and restoration drives a trade-oH in resistance to phages targeting the inner and outer core LPS

Phages Vale (CPL01276) and Tor (CPL00272) were selected to further investigate the resistance mechanisms and interactions of phages targeting different LPS regions. Tor, predicted to bind the outer core LPS, can infect the parent strain, which has complete LPS, but not Δ*pilA* Δ*galU,* which lacks the outer core LPS (Figure 2). Vale, predicted to bind the inner core LPS, can infect Δ*pilA* Δ*galU* but not the parent strain, where the inner core LPS is shielded by the outer core and O-antigen (Figure 1A and Figure 3).

Tor-resistant mutants of the parent strain were found to possess truncated LPS cores devoid of O-antigen, resembling the LPS observed in naïve Δ*pilA* Δ*galU* (Figure 4A). Vale could infect all three Tor-resistant mutants, with no significant difference in PFU relative to infecting naïve Δ*pilA* Δ*galU* (log10(PFU +1∼strain; Tor_R1: *p* = 0.662, Tor_R2: *p =* 0.949, Tor_R3: *p* =0.399, df = 24, SE = 0.1; Figure 4B).

**Figure 4:**
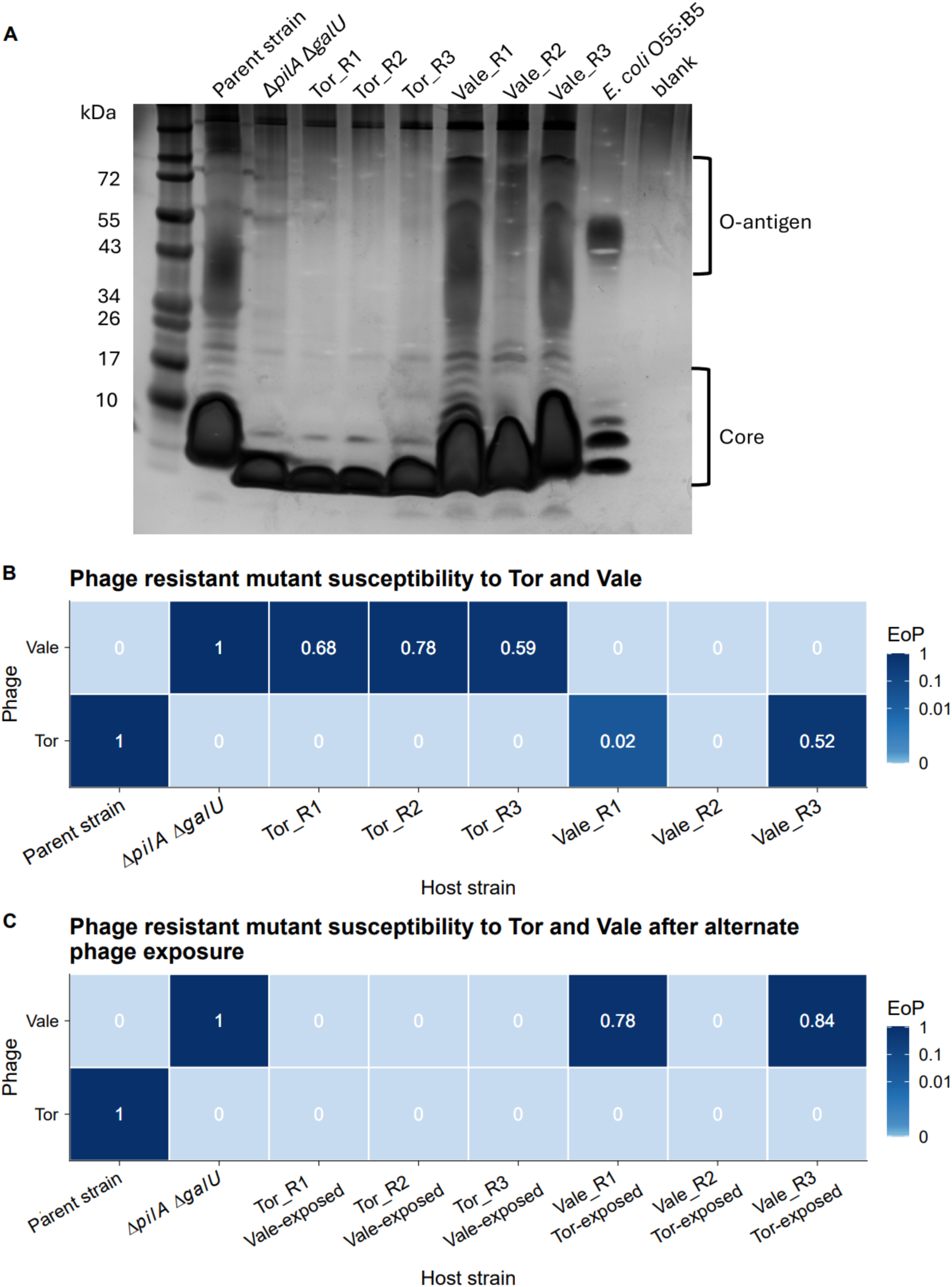
**[A]** The LPS extracted from the parent strain, Δ*pilA* Δ*galU,* three Tor-resistant parent strain mutants and three Vale-resistant Δ*pilA* Δ*galU* mutants, run on a 4-12% bis-tris SDS-PAGE gel. All three Tor-resistant parent strain mutants possess a truncated LPS core, devoid of O-antigen. Vale-resistant Δ*pilA* Δ*galU* have a restored LPS core, with O-antigen also restored in Vale_R1 and Vale_R3. **[B]** The mean efficiency of plating (EoP) of Tor and Vale on the parent strain, Δ*pilA* Δ*galU* and Tor- and Vale-resistant mutants. Resistance to Tor conferred susceptibility to Vale in all three mutants and resistance to Vale conferred susceptibility to Tor in two out of three mutants (Vale_R1 and Vale_R3). **[C]** The mean EoP of Tor and Vale on the phage resistant mutants after exposure to the alternate phage. All three Tor -resistant mutants retained resistance to Tor and gained resistance to Vale. Vale_R2 maintained resistance to both phages. Vale_R1 and Vale_R3 lost resistance to Vale and gained resistance to Tor.

All three Vale-resistant Δ*pilA* Δ*galU* mutants possessed larger LPS cores than naïve Δ*pilA* Δ*galU*, restored to similar sizes to those observed in the naïve parent strain (Figure 4A). Two of the Vale-resistant mutants, Vale_R1 and Vale_R3, also regained O-antigen. Thus, *P. aeruginosa* was capable of restoring wildtype LPS and O-antigen structures in response to phage predation through an unknown mechanism, as these strains lack the UDP-glucose synthesis pathway provided by *galU*. Restoration of outer core LPS and O-antigen in Vale_R1 and Vale_R3 conferred susceptibility to Tor, albeit with a significant 45.8-fold lower PFU relative to infection of the naïve parent strain for Vale_R1, whilst PFU on Vale_R3 was similar to infection of the naïve parent strain (log10(PFU +1)∼strain; Vale_R1: *p* < 0.0001, Vale_R3: *p* = 0.525, df = 24, SE = 0.161; Figure 4B). Vale_R2, with its partly restored LPS, was resistant to both Vale and Tor.

When Tor-resistant mutants were subsequently exposed to Vale, all three mutants maintained Tor resistance while also gaining resistance to Vale (Figure 4C). However, when Vale-resistant mutants (Vale_R1 and Vale_R3) were exposed to Tor, they gained resistance to Tor, but became resensitised to Vale, with no significant difference in PFU relative to infecting naïve Δ*pilA* Δ*galU* (log10(PFU +1) ∼ strain; Vale_R1: *p* =0.182, Vale_R3: *p* = 0.469, df = 16, SE = 0.0475; Figure 4C). Vale_R2, which was already resistant to both Vale and Tor, maintained resistance to both phages when exposed to Tor instead of Vale.

In summary, the parent strain gained resistance to Tor via LPS truncation and loss of O-antigen, conferring susceptibility to Vale. Meanwhile, Δ*pilA* Δ*galU* acquired resistance to Vale by restoring the outer core and O-antigen, conferring susceptibility to Tor in two out of three repeats. Subsequently exposing the phage-resistant mutants to the alternate phage revealed that Tor resistance was maintained but Vale-resistance was transient.

### Comparative genomics of phage resistant mutants reveals long deletions and SNPs potentially underpinning LPS modifications

Genomes of Tor- and Vale-resistant mutants were compared to their respective parent strains to identify SNPs, insertions and deletions relating to phage resistance (Figure 5A, Supplementary Table 4). Large deletions ranging from 100 kb to over 200 kb were identified in all three Tor-resistant mutants^36,41–44^ (Figure 5B). All three deletions included *galU*, involved in outer core LPS synthesis^45^, and *mexXY,* encoding an antibiotic efflux pump^36^. As these deletions affected *mexXY,* we tested whether resistance to Tor had increased aminoglycoside sensitivity. All three Tor resistant mutants were more susceptible to gentamycin than their naïve parent strain, with an minimum inhibitory concentration (MIC) of 0.25 µg/mL for Tor_R1 and Tor_R3 and 0.19 µg/mL for Tor_R2, compared 3 µg/mL to for the parent strain (Supplementary Figure 4). Tor_R1 and Tor_R3 also exhibited a red-brown phenotype (Supplementary Figure 5) and their deletions encompassed *hmgA,* which converts red homogentisic acid to colourless 4-maleylacetoacetate^36,44^. In Tor_R2, SNP analysis of short-read sequences detected a non-synonymous substitution (G11D) in *hcnB* (*pa2194*), encoding hydrogen cyanide synthase (Supplementary Table 5). *hcnB* is located at the edge of the long, deleted region, suggesting that the detected substitution is a by-product of the deletion rather than a separate adaptive mutation.

**Figure 5:**
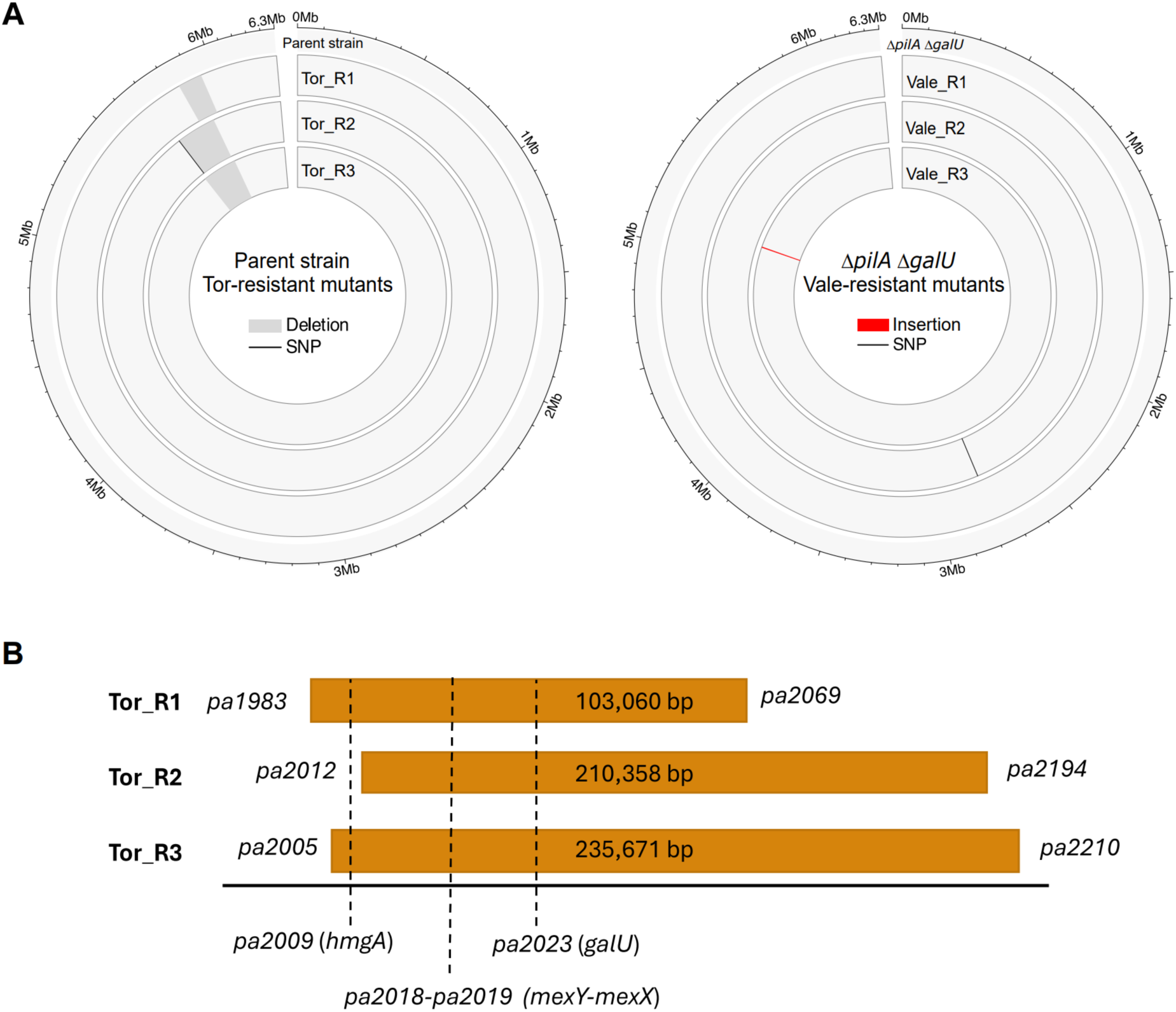
**[A]** Circos plots showing the SNPs, insertions and deletions in Tor- and Vale-resistant mutants relative to the parent strain and Δ*pilA* Δ*galU,* respectively. **[B]** Summary of the genes affected by the long deletions in Tor-resistant mutants.

A ∼7 kb insertion was detected in Vale_R3. Closer inspection revealed that this was pEX19gm-Δ*galU*, the plasmid used to generate the *galU* deletion, positioned upstream of the intact *galU* gene (Supplementary Figure 6). Sequence analysis and PCR amplification of *galU* identified no such insertion in Vale_R1 and Vale_R2 nor in their parental strain Δ*pilA* Δ*galU* (Supplementary Figure 6). In Vale_R2, SNP analysis revealed a substitution (P282S) in *wapH* (*pa5004*), which encodes a putative glycosyltransferase that is proposed to transfer D-glucose residues to the outer core^46,47^. GlycanInsight predicted the proline residue to be a part of the WapH binding pocket with a carbohydrate binding probability of 0.612^48^.

Long read sequencing of the phage-resistant mutants after exposure to the alternate phage showed that the long deletions in Tor-resistant mutants were maintained after exposure to Vale. No additional insertions or deletions were detected in the Tor-resistant mutants after Vale exposure. No mutations were detected in the Vale-resistant Δ*pilA* Δ*galU* mutants that were subsequently exposed to Tor.

### Combined treatment with phages targeting the LPS inner and outer core supresses resistance for at least 72 hours in PAO1

Growth curves were recorded over 72 hours for the parent strain and Δ*pilA* Δ*galU* treated with Tor and Vale, both independently and in combination (Figure 6). As expected, no differences were observed between Vale-treated parent strain, and Tor-treated Δ*pilA* Δ*galU*, compared to their respective untreated hosts, as these phages do not infect these naïve hosts. Phage resistance emerged after 20-24 hours for both the parent strain treated with Tor and Δ*pilA* Δ*galU* treated with Vale. The optical density of the Tor-treated parent strain exceeded that of the untreated parent strain, likely due to accumulation of red-brown pigment produced by Tor-resistant mutants (Supplementary Figure 5). Combined treatment with Tor and Vale suppressed growth in both the parent strain and Δ*pilA* Δ*galU* growth for the entire 72 hours. Thus, combined treatment with Tor and Vale, targeting different LPS regions, suppresses resistance for at least three-fold longer than each phage independently.

**Figure 6:**
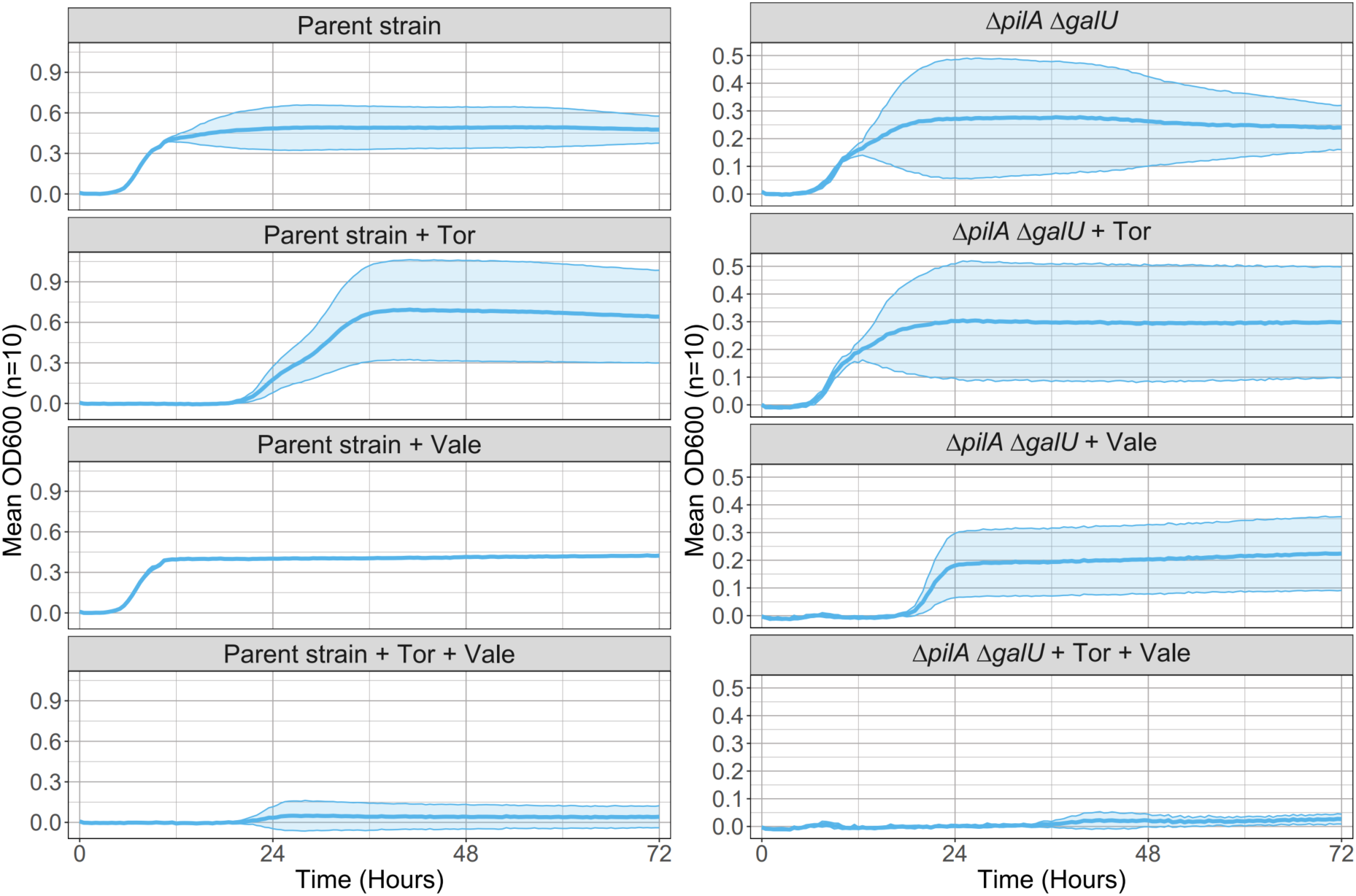
The growth curves of the parent strain and Δ*pilA* Δ*galU* in the presence of Tor and Vale separately and simultaneously. Ribbons represent the 95% confidence interval around the mean.

## Discussion

Reducing the likelihood of emergent phage resistance is crucial in designing effective phage cocktails. One strategy is to target a variety of phage receptors, or different structural morphologies of the same receptor, to suppress resistance via receptor modification ^22–24^. Here, we screened 50 *P. aeruginosa* phages from the CPL against a panel of receptor deletion mutants and found that they all require either the T4P, outer core LPS or O-antigen as their receptor. Enriching sewage samples on Δ*pilA* Δ*galU,* which lacks these three receptors, yielded twelve phages, which must utilise different receptors to the existing characterised phages in the CPL. We had aimed to increase the chance of obtaining OprM-targeting phages^18^ by enriching samples at a sublethal concentration of trimethoprim, as OprM can become hyper-expressed in response to antibiotics^49,50, 51^. However, all the novel phages could infect Δ*oprM,* indicating that none required OprM as a receptor. One of the novel phages, CPL01271, could infect all the deletion mutants, and while phage resistant mutants were recovered, its receptor remains unknown as no SNPs were identified from resistant mutant short-read sequences. CPL01271-resistant mutants were either phenotypically resistant^52^, with epigenetic or post-transcriptional modifications conferring resistance, or genetic modifications were present but not detected from short-read SNP analysis alone.

CPL01284 did not consistently form zones of lysis on LPS mutants but did proliferate on Δ*pilA ΔgalU* in liquid media, suggesting a differential ability to infect LPS mutants in liquid and solid media. Similar discrepancies in phage host range across solid and liquid media have been reported previously^53^. CPL01279S could not infect *ΔalgC* mutants, indicating that it may require the pel, psl or alginate exopolysaccharide as its receptor^54^. Phages Knedl and Clew-1 have previously been shown to bind psl exopolysaccharide^55,56^; Clew-1 was shown to effectively disrupt biofilms as psl is a crucial biofilm matrix component^57^. CPL01277, CPL01278 and CPL01275 exhibited inhibited infection on Δ*wbpL* and Δ*algC* mutants. Inability to form zones of lysis on Δ*wbpL* and Δ*algC* may be related to growth dynamics; Δ*algC* grows more slowly than the parent strain and Δ*wbpL* suffers a population decline following logarithmic growth, potentially related to its increased sensitivity to self-produced pyocins^58^. Finally, Vale, CPL01282, CPL1280 and CPL1279L could only infect strains with truncated LPS, indicating that the outer core LPS or O-antigen blocks these phages from accessing their receptor. Therefore, a likely receptor for these phages is a part of the LPS inner core, which is only exposed when the outer core or O-antigen is removed^26^.

Phages targeting the inner core LPS are clinically valuable as they can infect rough phenotype strains with truncated LPS, that more common outer core and O-antigen specific phages cannot infect. *P. aeruginosa* isolates from chronic cystic fibrosis lung infections frequently exhibit a rough phenotype, devoid of O-antigen^59,60^ and red mutants with truncated LPS, similar to the Tor-resistant mutants reported here, have previously been isolated from patients^61^. Furthermore, phages that bind the inner core LPS may be powerful additions to phage cocktails by infecting the rough phage resistant mutants that emerge following exposure to other LPS-targeting phages. Previously, an inner core-binding phage, PhiYY, was found to infect mutants resistant to the outer core-targeting phage PaoP5^26^. Similarly, we identified a trade-oM in resistance to Vale and Tor, binding the inner and outer core, respectively, mediated by LPS modifications. Resistance to Tor occurred via LPS truncation associated with long 100-200 kb deletions and conferred susceptibility to Vale. Furthermore, we observed that the inverse can also occur; resistance to Vale conferred susceptibility to Tor via restoration of the LPS and O-antigen in two of the three resistant mutants.

LPS truncation in Tor-resistant mutants could be attributed to 100 to over 200 kb deletions encompassing *galU,* required for UDP-glucose synthesis for a complete outer core^45^. These deletions also spanned *mexXY,* an antibiotic efflux pump and significant determinant of aminoglycoside resistance^36,62^. We observed increased gentamycin sensitivity in all three Tor-resistant mutants, with greater than 10-fold reduction in MIC. In two cases, *hmgA* was also deleted resulting in a red phenotype^36,44^. Similar red mutants with long deletions have been reported previously and associated with altered sensitivity to several antibiotics^36,41–44^, indicating a phage steering role for outer core LPS-targeting phages. The long deletions were maintained in the genome when Tor-resistant mutants were subsequently exposed to Vale, suggesting that the resistance afforded by the long genomic deletions and LPS truncation was stable and not readily reversed to provide resistance to Vale. It is unsurprising that these long deletions did not arise when Vale-resistant Δ*pilA* Δ*galU* mutants were subsequently exposed to Tor, as these mutants already lack the *galU* gene, so further deletion in this region would not lead to LPS truncation; Tor-resistance in the Vale-resistant Δ*pilA* Δ*galU* background must be arising via an alternative mechanism than LPS truncation derived from *galU* deletion.

Vale_R3 was found to be a merodiploid from the *galU* deletion process, with pEX19gm-Δ*galU* integrated upstream of the intact *galU* gene. Thus, Vale_R3 still expresses *galU,* allowing full LPS production. The finding suggests that a small proportion of the Δ*pilA* Δ*galU* mutant population were merodiploids that escaped sucrose counter-selection and only proliferated to detectable levels under phage selection. We found no evidence that Vale_R1 and Vale_R2 harboured intact *galU* genes from PCR or sequence data. It is surprising that Vale_R1 and Vale_R2 could restore their LPS core in the absence of *galU,* as *galU* is considered essential for UDP-glucose synthesis for a complete LPS core^45,63^. The substitution of a proline for a serine was detected in the binding pocket of WapH in Vale_R2, which expressed an intermediate-sized LPS with a restored outer core but no O-antigen. WapH is a putative glycosyltransferase proposed to add a glucose residue (GlcII) to the outer core by catalysing a 1,4-glycosisdic bond between UDP-glucose and the N-(alanyl)-galactosamine residue of the LPS core^46,47^. The mutation in the WapH binding pocket may alter its substrate specificity, allowing a different sugar donor than UDP-glucose to complete the outer core. This theory is consistent with the finding that Vale_R2 was resistant to both Vale and Tor; the altered outer core structure in Vale_R2 may be sufficient to block Vale from accessing the inner core, but lack the specific outer core used by Tor. In contrast, Vale_R1 has a complete outer core with O-antigen and is sensitive to Tor. However, the mechanism by which Vale_R1 restores its outer core remains unknown, as no mutations were identified by sequence analysis. As with the CPL01271-resistant mutants, the change in LPS phenotype may be epigenetically or post-transcriptionally controlled, or genetic modifications may have gone undetected using our methods. Mass spectrometry analysis of the Vale-resistant mutant LPS structures would be required to determine precisely how their cores are being completed in the absence of *galU*.

As resistance to one phage sensitises bacteria to the other, combined treatment with Tor and Vale suppressed resistance for at least 72 hours, compared to 20-24 hours for single phage treatments. This aligns with previous findings that phages binding the inner core LPS are powerful additions to phage cocktails with outer core or O-antigen targeting phages^23,26^. While growth of bacterial mutants resistant to both Tor and Vale was not observed in 72-hours of simultaneous phage treatment, sequentially exposing bacteria to Tor then Vale generated mutants resistant to both phages. This suggests that combined treatment with Tor and Vale may be more effective when applied simultaneously rather than sequentially. Previous studies have also found that simultaneous phage application results in quantitatively weaker phage resistance, with reduced likelihood of acquiring multiple resistance mutations^64,65^. The intermediate, cross-resistant LPS phenotype observed in Vale_R2 represents a likely mechanism of resistance that may emerge against simultaneous Tor-Vale treatment. Identifying a phage which infects Vale_R2 would further improve the depth of the Tor-Vale cocktail. A previous study found ‘white’ mutants that were resistant to both PhiYY and PaoP5, targeting the inner core and O-antigen, respectively^26^. PaoP5 was trained to produce PaoP5-m1 which could infect the ‘white’ mutants; a five-phage cocktail including PhiYY and PaoP5-m1 suppressed phage resistance for up to five days. Tor could be similarly trained to infect Vale_R2, or Vale_R2 could be used as an enrichment host to isolate phages capable of infecting this strain.

## Conclusions

Overall, this study provides an example of how strategically designing phage cocktails based on receptor specificity is crucial for minimising the likelihood of emergent phage resistance. Further investigation is required to determine whether a similar trade-oM between Tor and Vale would effectively suppress phage resistance in an *in vivo* infection model. Additionally, the selection landscape in a clinical setting and under antibiotic pressure may impact the emergence of phage resistant mutants with LPS modifications^59,60,66^. Tor and Vale could be combined with phages using different receptors to create a functionally diverse cocktail that is even less vulnerable to emergent resistance. The host deletion mutants engineered in this work provide both a valuable tool for identifying phage receptors in *P. aeruginosa,* and a powerful tool to further expand phage libraries to target receptors other than the LPS and T4P. In turn, this will assist in isolation and characterisation of novel phages for the strategic design of functionally diverse, personalised phage cocktails.

## Materials and methods

### Bacterial strains and growth conditions

The bacterial strains used in this study are listed in Table 1. Phages were provided by the CPL and are listed in the supplementary materials. Bacteria were grown in LB at 37 °C with 200 rpm shaking, unless stated otherwise. LB was supplemented with 10 mM MgCl_2_ and 10 mM CaCl_2_, herein referred to as sLB. Where specified, trimethoprim was added at 2.5 or 25 µg/mL. Phage-resistant mutants were always grown in the presence of phage to maintain selection pressure, with 20 µL of phage stock added per 10 mL of LB in liquid culture and 1 mL of phage stock added to 30 mL of LB agar to make solid plates.

### Generating a panel of unmarked receptor deletion mutants by homologous recombination

Unmarked deletions of common phage receptors were generated by two-step allelic exchange^67^ in the promiscuous *P. aeruginosa* PAO1 Δ*hsdR* strain^68^. ∼500bp regions flanking the target genes were amplified, joined together by overlap extension PCR and ligated into the plasmid pEX19gm. Modified plasmids were transformed into *Escherichia coli* DH5α for amplification, and then into *E. coli* Rho3 for conjugation into the PAO1 recipient strain on SOB agar (1.5% bacteriological agar, 20 mM bacto-tryptone, 5 mM bacto-yeast extract, 0.5 mM NaCl, 2.5 mM KCl, 10 mM MgCl_2_). Double recombinants were obtained by counter-selective plating on 100 µg/mL gentamycin and 5% sucrose. Successful mutants were iden9fied by PCR amplifica9on and Sanger sequencing of the ∼1000bp region surrounding the target gene. Δ*pilA* Δ*galU* and Δ*pilA* Δ*algC* mutants were constructed by first dele9ng *pilA* and then the respec9ve LPS synthesis gene. All dele9ons were performed on LB agar, except the dele9on of *wbpL,* which required MOPS-pyruvate minimal media (for 1 litre: 100 mL 10× MOPS (400 mM MOPS buffer, 40 mM tricine, 0.1 mM FeSO_4_, 95 mM NH_4_Cl, 2.76 mM K_2_SO_4_, 5 μM CaCl_2_ and 5.25 mM MgCl_2_, adjusted to pH 7.6 with ∼330 mM KOH), 10 mL 0.132 M K_2_HPO_4_, 10 mL 20% *w/v* sodium pyruvate, 15 g Bacto agar), as O-an9gen deficient strains are sensi9ve to self-produced pyocins on nutrient-rich media^58^.

### High-throughput receptor identification by spot assays on the deletion mutant panel

A 96-well microtitre plate (Grenier Bio-One, Austria) was set up with a random arrangement of 50 *P. aeruginosa* phages, 9 T7 positive controls and 29 sLB blanks without phage. Phage wells contained 1 mL of sLB, 5 µL of overnight host culture (*E. coli* BW or *P. aeruginosa* PAO1) and 5 µL of phage lysate. Following overnight incubation (37°C, 200 rpm) to enrich phage, 100 µL from each well was filtered by centrifugation at 900 × g for 4 minutes through a 0.45 µm pore filter MultiScreenHTS HV sterile filter plate (PVDF membrane, Millipore, Merck) to remove bacteria. Bacterial overlay plates were set up by mixing 2 mL of mid-logarithmic phase (OD_600_ = 0.6) bacterial culture with 6 mL of top agar (0.65% w/v bacteriological agar, 10 mM MgCl_2_, 10 mM CaCl_2_) and poured over a 60 mL bottom agar plate (1% w/v bacteriological agar, 10 mM MgCl_2_, 10 mM CaCl_2_)^69^. 2µl of filtrate from each well was spotted onto a bacterial lawn of each of the panel of deletion mutants and *E. coli* and incubated overnight at 37 °C, and zones of lysis observed the following day. To confirm there was no well-to-well contamination no plaques are expected to form from the blank wells and only wells containing T7 form plaques on *E. coli*.

### Phage isolation using *ΔpilA ΔgalU* as an enrichment host

Wastewater samples were provided to the CPL by the Environment Agency as 100 µL aliquots and stored at −20 °C. To obtain phages that use receptors other than T4P and outer core LPS, 95 wastewater samples were enriched on Δ*pilA* Δ*galU* and Δ*pilA* Δ*algC.* Another 95 wastewater samples were enriched on Δ*pilA* Δ*galU* with trimethoprim added to the growth media at 2.5 and 25 µg/mL, to encourage the upregulation of antibiotic resistance machinery. Phages were isolated and purified following the CPL enrichment workflow, reported previously^37^. Briefly, wastewater samples underwent two rounds of enrichment with the bacterial host and 0.22 µm-filtration. 2 µL of each enriched sample was spotted onto a host bacterial lawn and incubated overnight at 37 °C. Cores were picked from zones of lysis, transferred to 100 µl SM buffer (50 mM Tris-HCl (pH 7.5), 0.1 M NaCl, 8 mM MgSO_4_) and purified by two rounds of dilution to extinction.

### Phage DNA extraction, sequencing and assembly

50 µL of SM buffer containing a phage agar core and 500 µL of overnight bacterial host culture were added to two falcon tubes containing 20 mL of sLB and incubated overnight at 37 °C^37^. The two 20 mL cultures were combined into one and centrifuged at 10,000 × g for 30 minutes at 4 °C. The supernatant was filtered using a 0.22 µm syringe filter. 30 mL of filtrate was treated with 5 µg/mL DNase I (Roche, Merck, Darmstadt, Germany) and 10 µg/mL RNase A (Invitrogen, ThermoFisher Scientific, Waltham, MA, USA) for 30 minutes at 37 °C to remove bacterial nucleic acid. To precipitate phages, 10% *w/v* polyethylene glycol 8000 (PEG8000) and 1 M NaCl were added, mixed by inversion until dissolved, and incubated overnight at 4 °C. Precipitated phages were pelleted by centrifugation at 10,000 × g for 30 minutes and resuspended in 1 mL SM buffer. The DNA was extracted using the Norgen Phage DNA Isolation Kit (Norgen Biotek, Thorold, ON, Canada). Phage genomes were sequenced by the Exeter Sequencing Facility on an Illumina Novaseq SP platform with 2 x 150 bp paired-end reads.

Phage genomes were assembled and annotated as described in Tong *et al*. (2025)^68^. Briefly, reads were QCed with FASTP (v. 0.24.1) with the following settings (--dedup --dup_calc_accuracy 6 --length_required 30 –correction), and high-quality reads were mapped against the genome of the propagation host with minimap2 (v. 2.26) to remove any residual host DNA. Unmapped reads were subsampled to 500-fold coverage with shovill (v. 1.1.0) and assembled with unicycler (v. 0.5.0). Assembly graphs were manually inspected to confirm single circular genomes, with branches of up to 3 bp manually resolved by selection of the most abundant node. Annotation of phage genomes was performed with pharokka (v. 1.7.1) followed by phold (v. 0.1.3).

PhageClouds was used to identify phages form the NCBI genbank database that were closely related to the novel phages^70^. VIRIDIC was used to generate a plot of the novel phages, their closest relatives and other phages discussed in this study based on relatedness^71,72^.

### Preparation of phage-resistant mutants

Tor-resistant parent strain mutants and Vale-resistant Δ*pilA* Δ*galU* mutants were generated to investigate the resistance mechanisms of phages using the outer and inner LPS core, respectively. Five cultures of each host were grown from single colonies to OD_600_ of 0.6 in 25 mL of sLB, infected with 50 µL of the relevant phage lysate, and incubated overnight to allow phage-resistant mutants to emerge. Each culture was streaked out on 30 mL of LB agar containing 1 mL of phage lysate to obtain single resistant colonies. A single colony from each plate was used to inoculate 10 mL of sLB containing 20 µL phage lysate and grown to OD_600_ of 0.6. 1 mL of each culture was added to 500 µL glycerol to make a −80 °C cryostock. 1 mL of culture was also added to 3 mL top agar and poured over a 30 mL bottom agar plate. 5 µL of phage lysate was spotted onto the bacterial lawn, and after overnight incubation, a lack of clearance confirmed that the mutants were phage resistant.

Single colonies of three resistant mutants for each phage (Tor_R1, Tor_R2, Tor_R3, Vale_R1, Vale_R2 and Vale_R3), the naïve parent strain and naïve Δ*pilA* Δ*galU* were grown overnight in 10 mL LB, containing 20 µL phage lysate. The cultures were diluted 1:50 in 500 mL LB containing 1 mL phage lysate and incubated at 37 °C with 200 rpm shaking. When the culture reached an OD_600_ of 0.6, 2 mL of culture was added to 6 mL of top agar and poured over a 60 mL bottom agar plate for a spot assay to investigate cross-resistance between Tor and Vale. Tor and Vale lysates were serially diluted in LB to 10^-9^ with four repeats, and 5 µl of each dilution was spotted onto the bacterial lawn. After overnight incubation at 37 °C, plaques were enumerated to calculate plaque forming units (PFU) and efficiency of plating (EoP) relative to infection on the parent strain^69^. The remaining culture was grown to an OD_600_ of 1.0. 1 mL of culture was added to 500 µL glycerol to create another −80 °C cryostock and 1 mL of culture was aliquoted into a microcentrifuge tube for DNA extraction. The remaining culture was pelleted, 250 mL at a time, at 4,000 × g for 15 mins, to give ∼500 mg of cells for the LPS extraction.

The second round of phage resistant mutants were generated by growing single colonies of the original phage resistant mutants overnight in 10 mL LB with 20 µL of phage lysate of the alternate phage, such that Tor_R1, Tor_R2 and Tor_R3 were exposed to Vale phage lysate and Vale_R1, Vale_R2 and Vale_R3 were exposed to Tor phage lysate. As before, the cultures were streaked out onto LB agar containing 1 mL of phage lysate. Single colonies were picked and grown to OD_600_ 0.6 in 10 mL of sLB containing 20 µL phage lysate. 2 mL of culture was added to 6 mL of top agar and poured over a 60 mL bottom agar plate. Tor and Vale were serially diluted in LB to 10^-9^ with three repeats, and 2.5 µL of each dilution was spotted onto the bacterial lawn. After overnight incubation at 37°C, PFU and EoP were calculated.

Two CPL01271-resistant parent strain mutants were generated using the same method of infecting 30 mL cultures with phage and streaking them out onto LB agar containing phage. Two single colonies were selected and grown overnight in 10 mL LB containing 20 µL phage lysate. 1 mL of each culture was used to set up a spot assay to confirm resistance, as before, and 1 mL of culture was aliquoted into a microcentrifuge tube for DNA extraction.

### LPS extraction and analysis from phage resistant mutants

LPS was isolated using a method adapted from Darveau and Hancock^73^. The 500 mg cell pellets were resuspended in 15 mL 10 mM Tris-hydrochloride buffer (pH 8), containing 2 mM MgCl2_2_, 100 µg/mL DNase I and 25 µg/mL RNase A. The cells were sonicated using a Sonics VCX130PB sonicator at a frequency of 20 kHz with a 6 mm diameter probe for 16 30-second bursts at 75% probe intensity, such that a total of 8500 J was applied to the sample. DNase I and RNase A were added to final concentrations of 200 and 50 µg/mL, respectively, and incubated at 37 °C for two hours. 5 mL 0.5 M tetrasodium EDTA dissolved in 10 mM Tris-HCl and 2.5 mL of 20% SDS (each at pH 8) were added to the sample. The sample was vortexed and centrifuged at 50,000 × g at 20°C to remove peptidoglycan. Proteinase K was added to the supernatant at 4 units/mL and incubated overnight at 37 °C with shaking at 200 rpm. Precipitate was removed by centrifuging at 200 × g 10 min. Two volumes 0.375 M MgCl_2_ in 95% ethanol were mixed with the supernatant, cooled to 0 °C in a −20 °C freezer, and centrifuged at 12,000 × g for 15 minutes at 0 °C. The pellet was resuspended in 25 mL 2% SDS-0.1 M tetrasodium EDTA, dissolved in 10 mM Tris-HCl (pH8) and sonicated at 20 kHz with a 6 mm diameter probe for two 30s bursts at 75% probe intensity, applying at total of 1000 J to the sample. The sample was incubated at 85 °C for 20 minutes and proteinase K added at 0.5 units/mL and incubated overnight at 37 °C at 200 rpm to ensure complete denaturation of SDS-resistant proteins. Two volumes of 0.375 M MgCl_2_ in 95% ethanol were added, cooled to 0 °C, as measured with a thermometer in a −20 °C freezer, and centrifuged as before. The pellet was resuspended in 15 mL 10 mM Tris-HCl (pH 8) and sonicated again for two 30s bursts at 75% probe intensity. The sample was centrifuged at 200 × g for 5 min to remove insoluble Mg^2+^-EDTA crystals. The supernatant was centrifuged at 200,000 × g for 2 hours at 15 °C in the presence of 25 mM MgCl_2_. The pellet was resuspended in 100 µL of nuclease-free water.

5 µL of LPS sample was added to 5 µL Nu-PAGE LDS sample buffer (4x) and 10 µL nuclease free water and incubated at 100 °C for 10 mins. Samples were then loaded onto a 4-12% Bis-Tris SDS gel and run for 60 min at 150 V. Gels were stained using the Proteosilver Plus Silver Stain Kit.

### Resistant mutant DNA extraction, sequencing and comparative genomics

1 mL of resistant mutant culture was washed three times with LB to remove phage and pelleted at 10,000 × g for 5 minutes. DNA was extracted using the Promega Wizard DNA extraction kit. Libraries were prepared following the rapid sequencing DNA V14 - barcoding protocol (SQK-RBK114.24). Genomes were sequenced by the Exeter sequencing Facility using Illumina Novaseq SP for short-reads and an Oxford Nanopore PromethION R10 flowcell for long-reads with 2 × 150 bp paired-end reads. Tor- and Vale-resistant mutants underwent both long- and short-read sequencing, while only long reads were obtained for the second-round resistant mutants where Tor- and Vale-resistant mutants were exposed to the alternate phage. Only short reads were obtained for CPL01271-resistant mutants. Raw long reads were quality filtered, and adapters were removed with porechop (v. 0.2.4, https://github.com/rrwick/Porechop), discarding reads with internal adapters. Short-read only assemblies were performed using unicycler (v. 0.5.1, http://github.com/rrwick/Unicycler). Long-read assemblies were performed using Trycycler (v. 0.5.5, https://github.com/rrwick/Trycycler), followed by short-read polishing with polypolish (v. 0.6.0, https://github.com/rrwick/Polypolish). Genomes were annotated using prokka V1.14.6^74^. For Tor- and Vale-resistant strains, complete circular chromosomes were obtained. In both the parent strain and Δ*pilA* Δ*galU* groups, the genomic sequences were adjusted to start in the same position and strand orientation.

For CPL01271-resistant strains, Snippy (v 4.6.0, https://github.com/tseemann/snippy) was used with default parameters to detect SNPs. To detect large insertions and deletions in the resistant mutants, pairwise alignments of long-read sequenced genomes were performed using nucmer V4.0 (MUMmer package)^75^ with a minimum cluster length of 5000 bp. The coordinates of large deletions and insertions were exported using the show-coords command on the delta files. SNPs were detected in short-read sequenced genomes using Snippy with the default parameters. Deletions, insertions and SNPs bed files were merged for each genome. Circos plots for both groups were generated in R using the library circlize V 0.4.15^76^. The web-based tool GlycanInsight (https://www.glycaninsight.cn/index/), which uses the DeepGlycanSite algorithm to predict carbohydrate-binding probabilities, was used to predict whether the SNP found in LPS biosynthesis gene *wapH* occurred in the binding pocket^48^.

### Gentamycin susceptibility assay

Several colonies were taken from a fresh streak plate and resuspended in 0.85% saline solution to McFarland standard 0.5. The culture was spread across a Mueller-Hinton plate (1.5% w/v bacteriological agar) using a sterile swab to achieve an even bacterial lawn. A Liofilchem gentamycin MIC test strip was placed on the plate using sterile forceps. After overnight incubation at 37 °C, the zone of clearance was measured against the test strip to determine the MIC of gentamycin on each strain.

### Growth curve assays

Growth curves were generated over 72 hours for the parent strain and Δ*pilA* Δ*galU* with separate and combined treatments of Tor and Vale, and in the absence of phage. 10 mL of LB was inoculated with several colonies from fresh streak plates, grown to OD_600_ of 0.6 (37 °C, 200rpm), and then diluted to OD_600_ of 0.1. A 96-well plate was set up with six replicates of each bacterial control well (95 µL sLB, 5µL bacterial culture), phage control well (98 µL sLB, 2 µL phage), single phage treatment well (93 µL sLB, 5 µL bacterial culture, 2 µL phage at MOI 0.1), and double phage treatment well (91 µL sLB, 5 µL bacterial culture, 2 µL of each phage at MOI 0.1).

Growth curves were also generated for each deletion mutant in the absence of phage to determine whether the deletions had cause fitness defects. 10 mL of LB was inoculated with colonies from fresh streak plates and incubated for 3 hours (37 °C, 200rpm). Cultures were diluted to a starting OD_600_ of 0.1 and 100 µL of each culture was added to six replicate wells of a 96-well plate. Optical density was recorded every 15 minutes for 16 hours in a plate reader (Tecan Sunrise). The intrinsic growth rate (r), carrying capacity (K) and doubling time (DT) were calculated using Growthcurver^77^.

### Statistical analysis

A linear model was used to determine whether the log-transformed mean PFU significantly differed between resistant mutants and their parent strain (log10(PFU + 1) ∼ strain). The estimated marginal means were extracted from the model using the emmeans package (https://CRAN.R-project.org/package=emmeans). Linear models were used to determine whether mean intrinsic growth rate, carrying capacity and doubling time differed significantly between deletion mutants and wildtype PAO1 (growth parameter∼strain).

## Abbreviations

LPS: Lipopolysaccharide
T4P: Type IV Pili
SNP: Single Nucleotide Polymorphism

## Data availability

Source data and information on the phages used in this study are provided in the supplementary materials. NCBI Accession numbers for phages used in this study are provided in the Supplementary materials. Sanger sequences of the deletion mutants are available at: https://github.com/citizenphage/Tong-272-1276.

## Code availability

The data analysis in this study was performed with R and python code, available at https://github.com/citizenphage/Tong-272-1276.

## Acknowledgments

The authors would like to acknowledge the contributions of Tiandi Yang and Anne Dell for their advice and support in LPS extraction and analysis.

## Author contributions

ET constructed the panel of deletion mutants, performed the experiments and wrote the manuscript. BT and ET conceived the experimental design. LB, BT and ET performed phage and resistant mutant genome assemblies and alignments. ET, JF, RM, CF, CB, JS and JWS isolated and purified phages from water samples collected by citizen scientists: RA, JG, NW, and TI. BT and SP edited the manuscript.

