## Supplementary figures and images for "Lipopolysaccharide truncation and restoration drives a trade-off in resistance to two phages in *Pseudomonas aeruginosa*"

### Supplementary Figure 1

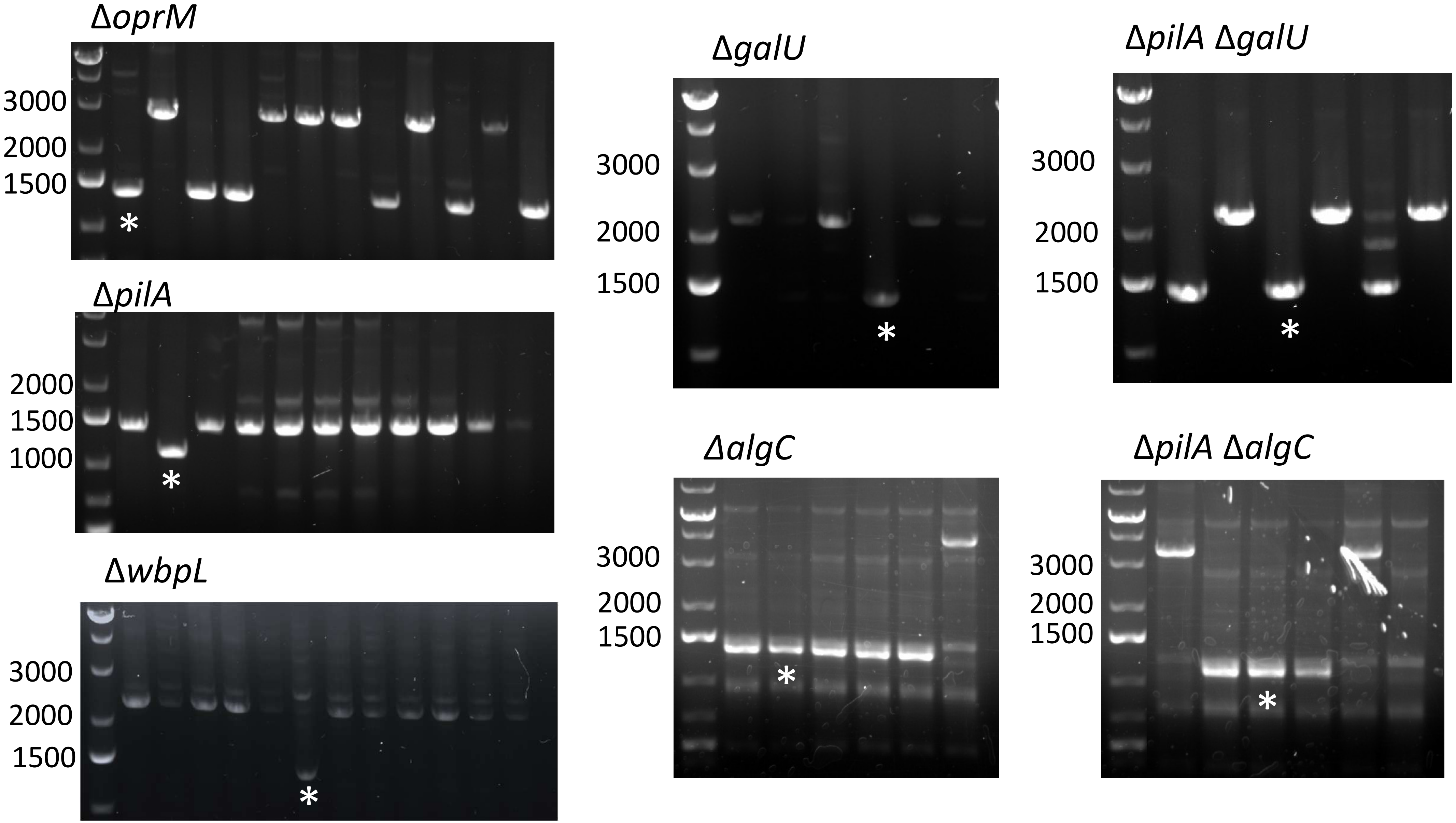

### Supplementary Figure 2

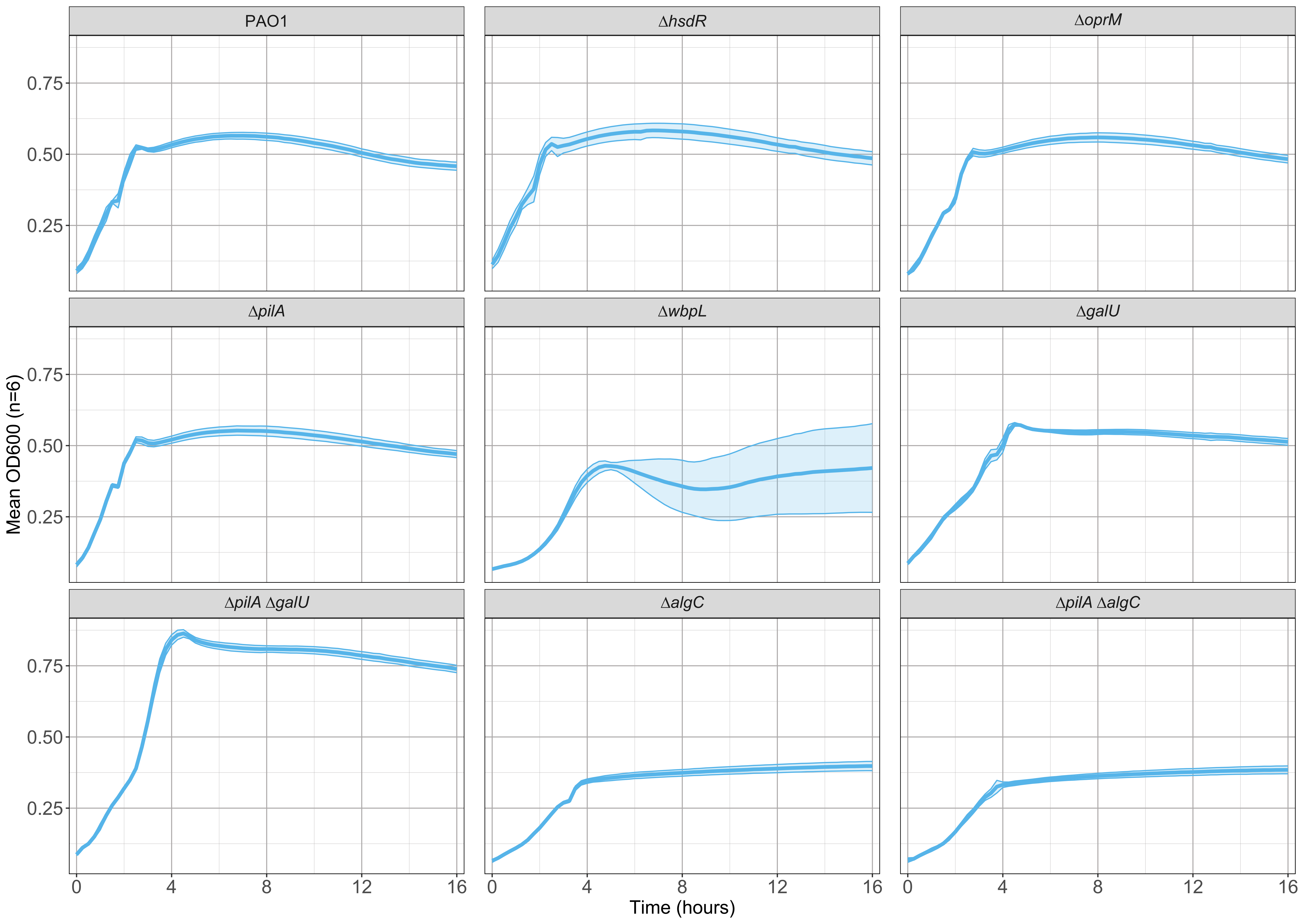

### Supplementary Figure 3

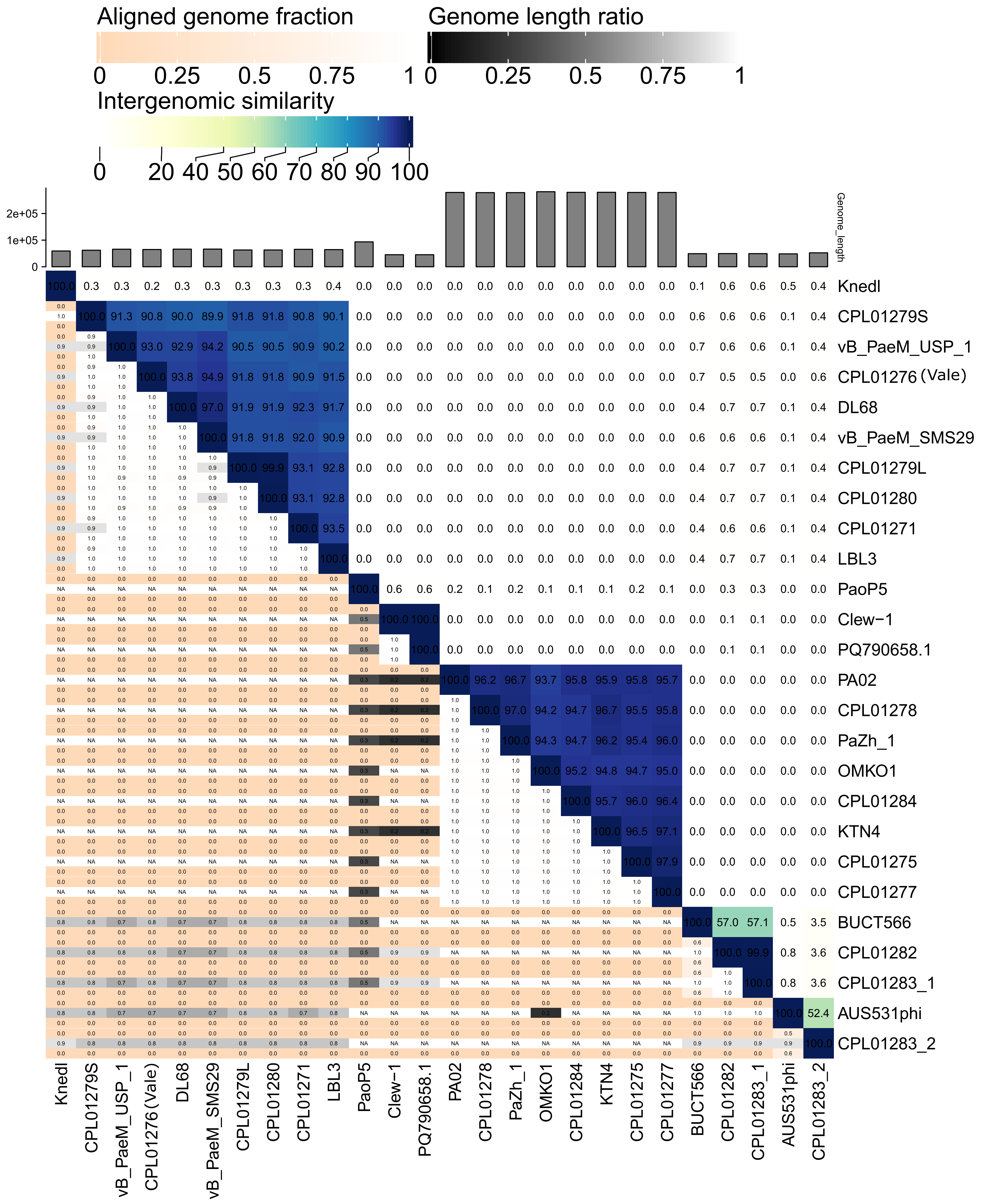

### Supplementary Figure 4

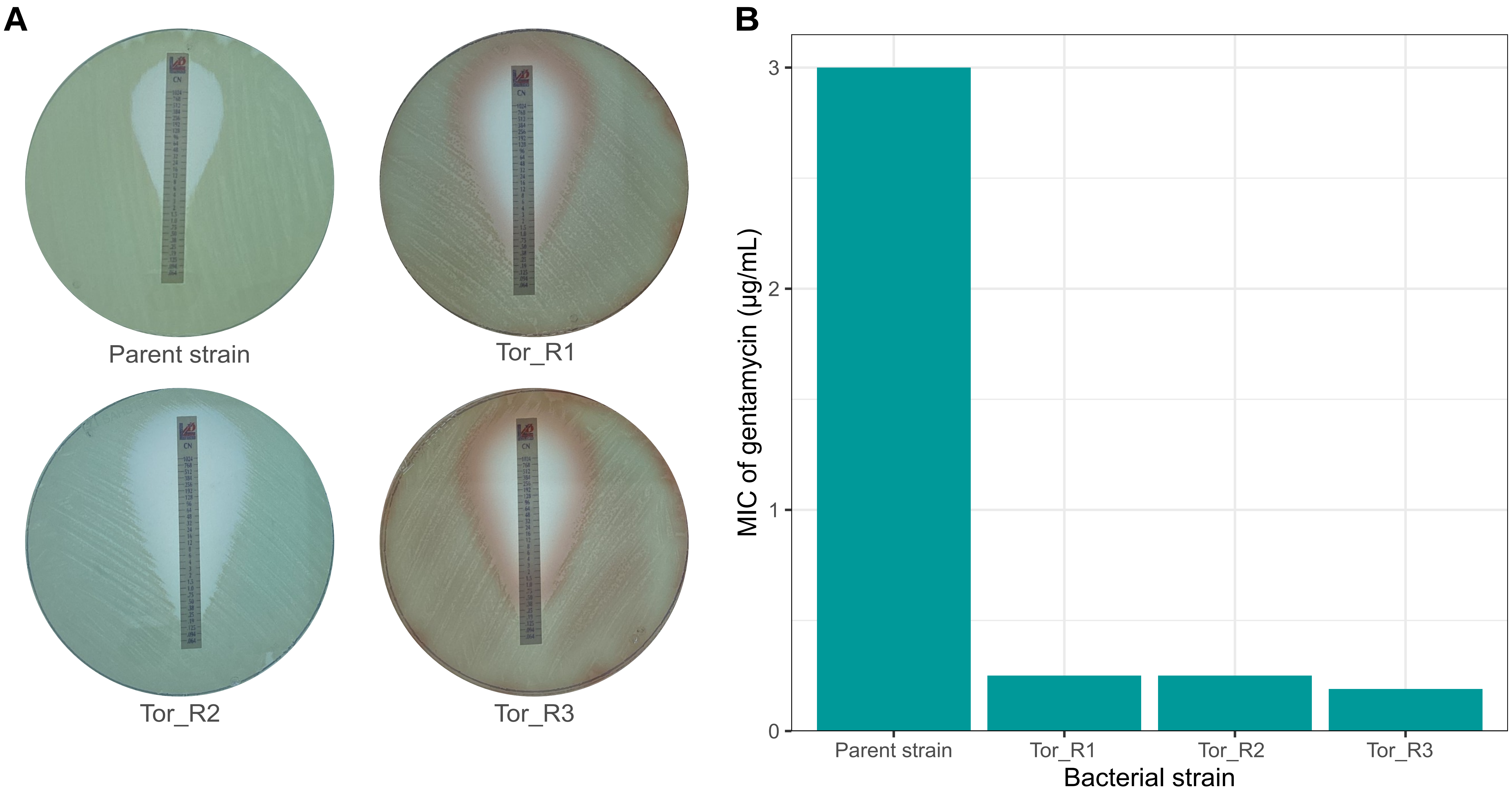

### Supplementary Figure 5

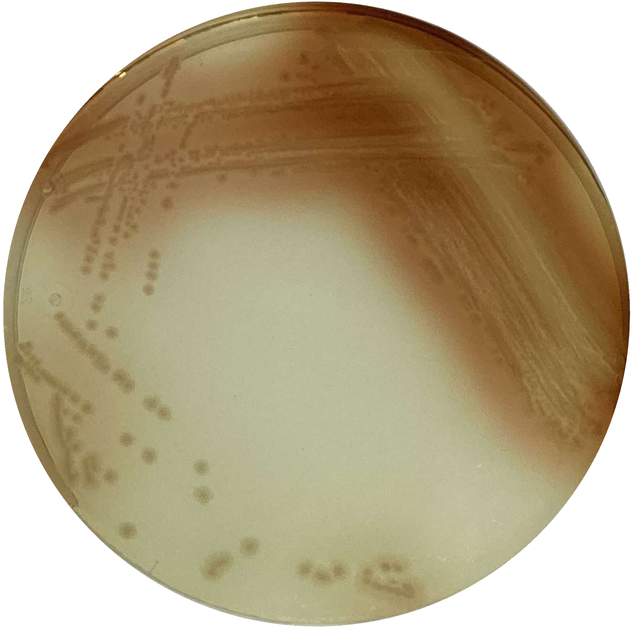

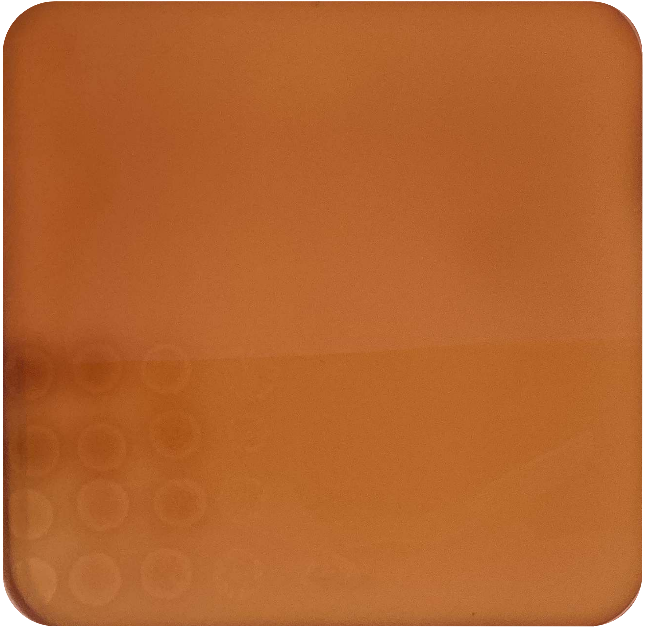
