## Supplementary Figure 6 for "Lipopolysaccharide truncation and restoration drives a trade-off in resistance to two phages in *Pseudomonas aeruginosa*"

**A**

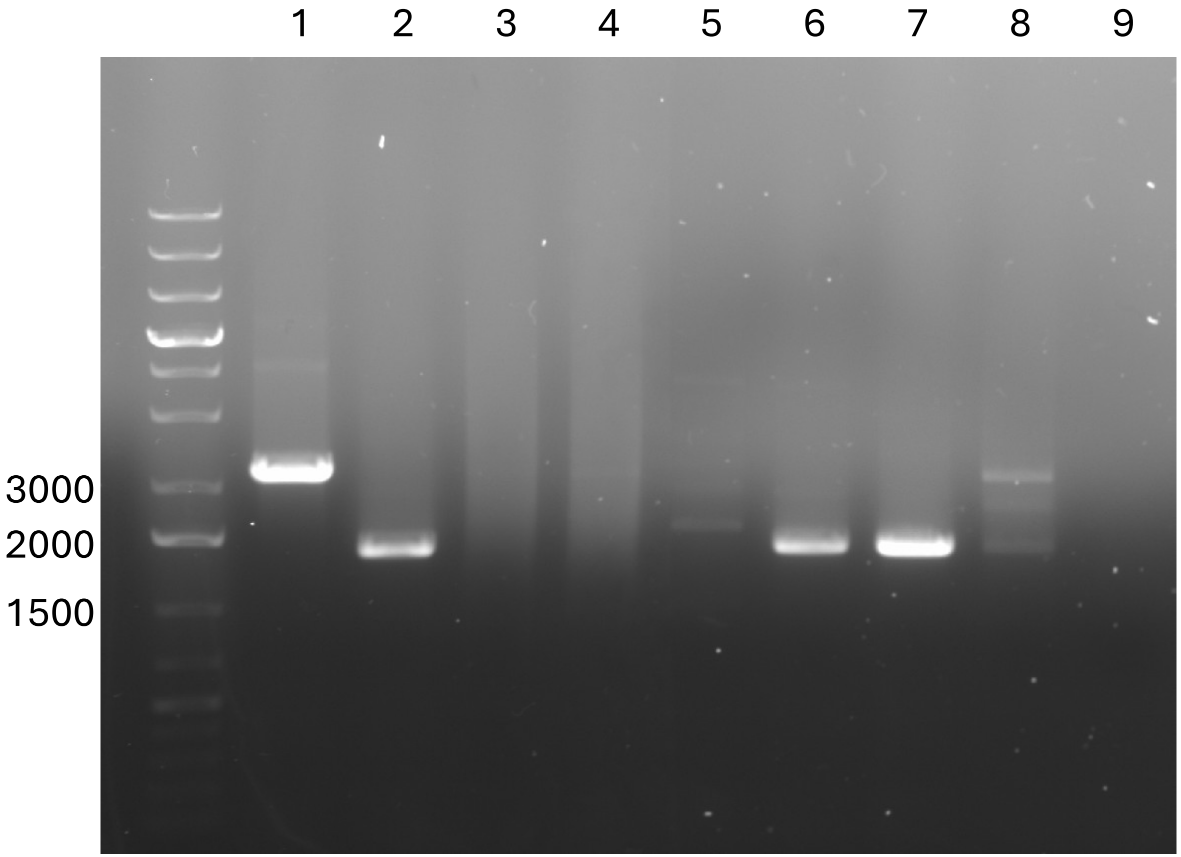


1. Parent strain (complete LPS)
2. Δ*pilA* Δ*galU* (outer core to O-antigen deletion)
3. Parent strain Tor_R1
4. Parent strain Tor_R2
5. Parent strain Tor_R3
6. Δ*pilA* Δ*galU* Vale_R1
7. Δ*pilA* Δ*galU* Vale_R2
8. Δ*pilA* Δ*galU* Vale_R3
9. NFW blank

**B**

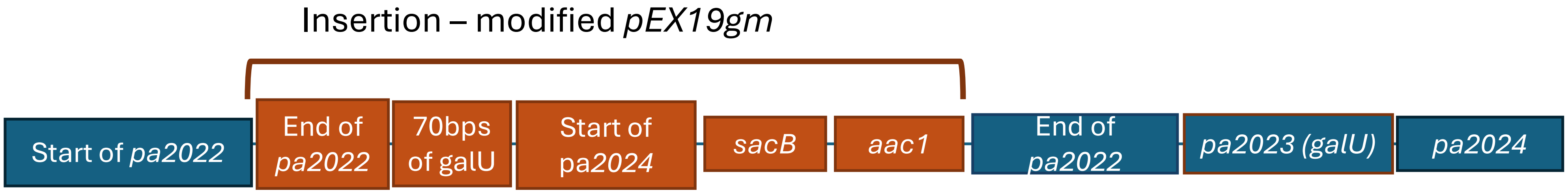
