## Supplementary Materials for "Lipopolysaccharide truncation and restoration drives a trade-off in resistance to two phages in *Pseudomonas aeruginosa*"

**Supplementary Material**

| **Plasmid** | **Description** | **Accession no.** | **Source** |
| --- | --- | --- | --- |
| pEX19gm | Allelic exchange vector encoding gentamycin resistance (*aaC1*) and sucrose sensitivity (sacB) | KM887142 | Hoang et al (1998) ^1^ |
| pEX19gm-∆*oprM* | Modified pEX19gm containing ~500 bp regions that flank *oprM* |  | This study |
| pEX19gm-∆*pilA* | Modified pEX19gm containing ~500 bp regions that flank *pilA* |  | This study |
| pEX19gm-∆*wbpL* | Modified pEX19gm containing ~500 bp regions that flank *wbpL* |  | This study |
| pEX19gm-∆*galU* | Modified pEX19gm containing ~500 bp regions that flank *galU* |  | This study |
| pEX19gm-∆*algC* | Modified pEX19gm containing ~500 bp regions that flank *algC* |  | This study |

**Supplementary Table 1:** The plasmids used to construct the panel of knockout mutants.

| **Primer** | **Sequence** | **Description** |
| --- | --- | --- |
| PA0427P1 | CGCGGATCCCCGAAAGGCGTTGGCTACTC | Forward primer for *oprM* upstream region and overlap extension |
| PA0427P2 | GGGATCTTCCTTCTTCGCGGTGGAAAGGAAGGACCGTTTC | Reverse primer for *oprM* upstream region |
| PA0427P3 | GAAACGGTCCTTCCTTTCCACCGCGAAGAAGGAAGATCCC | Forward primer for *oprM* downstream region |
| PA0427P4 | ACACGAATTCGACAATTTCGGCAACCGCGC | Reverse primer for *oprM* downstream region and overlap extension PCR |
| PA0427checkF | AAGCTGGAGCGCTACAATGG | Forward primer for *oprM* check PCR |
| PA0427checkR | GAAGTCGACGGTAATCGCGATC | Reverse primer for *oprM* check PCR |
| PA4525P1 | ACGCGGTACCAAGCCGGTGGAAGTGGAAGTG | Forward primer for *pilA* upstream region and overlap extension |
| PA4525P2 | GAACTGATGATCGTGGTTGCGATCCTGAACCGTACTGCGGATG | Reverse primer for *pilA* upstream region |
| PA4525P3 | CATCCGCAGTACGGTTCAGGATCGCAACCACGATCATCAGTTC | Forward primer for *pilA* downstream region |
| PA4525P4 | ACACGAATTCCAGAGGGATGACCCGGTGTTG | Reverse primer for *pilA* downstream region and overlap extension PCR |
| PA4525checkF | ACCATCGCATCGGCCTCTAC | Forward primer for *pilA* check PCR |
| PA4525checkR | CAGAATCGCTTCGGTCGTCAG | Reverse primer for *pilA* check PCR |
| PA3145P1 | CGCGGATCCGTGCAGGCGGTGATAGCCAAAGTG | Forward primer for *wbpL* upstream region and overlap extension |
| PA3145P2 | GGATGATCGCGTGTCTAGTTCTCTTGGCGGTAGGATACAAGGC | Reverse primer for *wbpL* upstream region |
| PA3145P3 | GCCTTGTATCCTACCGCCAAGAGAACTAGACACGCGATCATCC | Forward primer for *wbpL* downstream region |
| PA3145P4 | ACACGAATTCGTATGGAGTTGGTGGTTGTCC | Reverse primer for *wbpL* downstream region and overlap extension PCR |
| PA3145checkF | AAGCCGACGGCTCAGTATAG | Forward primer for *wbpL* check PCR |
| PA3145checkR | GTAACAGCACCCGGCAACAG | Reverse primer for *wbpL* check PCR |
| PA2023P1 | ACACAAGCTTGGGCTACGACAGCCAGATAC | Forward primer for *galU* upstream region and overlap extension |
| PA2023P2 | CAGTGAGCCTTGCCGGTCTTGAAACGGGTGCCGTAAC | Reverse primer for *galU* upstream region |
| PA2023P3 | GTTACGGCACCCGTTTCAAGACCGGCAAGGCTCACTG | Forward primer for *galU* downstream region |
| PA2023P4 | ACACGAATTCCTACGAAACCTTCTGCGGCATG | Reverse primer for *galU* downstream region and overlap extension PCR |
| PA2023checkF | AGCGCATCGGTACCCATTTC | Forward primer for *galU* check PCR |
| PA2023checkR | GATGGGCTATGCTTGTCCGTG | Reverse primer for *galU* check PCR |
| PA5322P1 | CGCGGATCCGACCGCCTCAACCAGCAATG | Forward primer for *algC* upstream region and overlap extension |
| PA5322P2 | CAGTTGGTTACGGAAGACGGTCAACTCGTTCCCATCCTTG | Reverse primer for *algC* upstream region |
| PA5322P3 | CAAGGATGGGAACGAGTTGACCGTCTTCCGTAACCAACTG | Forward primer for *algC* downstream region |
| PA5322P4 | ACACGAATTCGCATGTTCAGCAGCCCGACGTTG | Reverse primer for *algC* downstream region and overlap extension PCR |
| PA5322checkF | ACAGCGACGAGAACGCGATCACG | Forward primer for *algC* check PCR |
| PA5322checkR | GATGTTGTAGGACTCGCCG | Reverse primer for *algC* check PCR |

**Supplementary Table 2:** The primers used to construct the panel of knockout mutants.

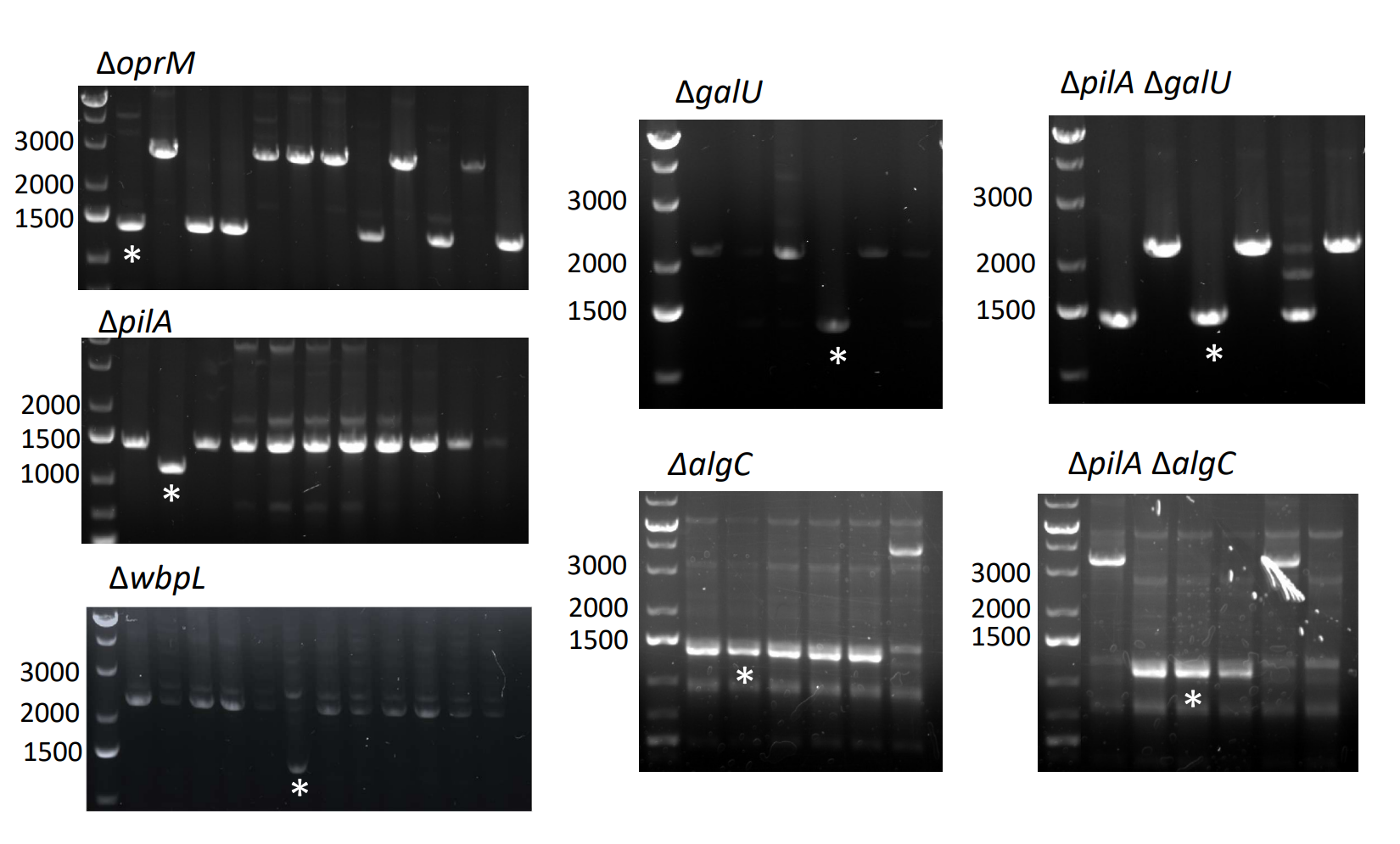

**Supplementary Figure 1:** The check PCRs confirming successful deletions for the panel of receptor knockouts. Successful in-frame deletion bands (*) were excised, gel extracted and Sanger sequenced to confirm the clean deletions.

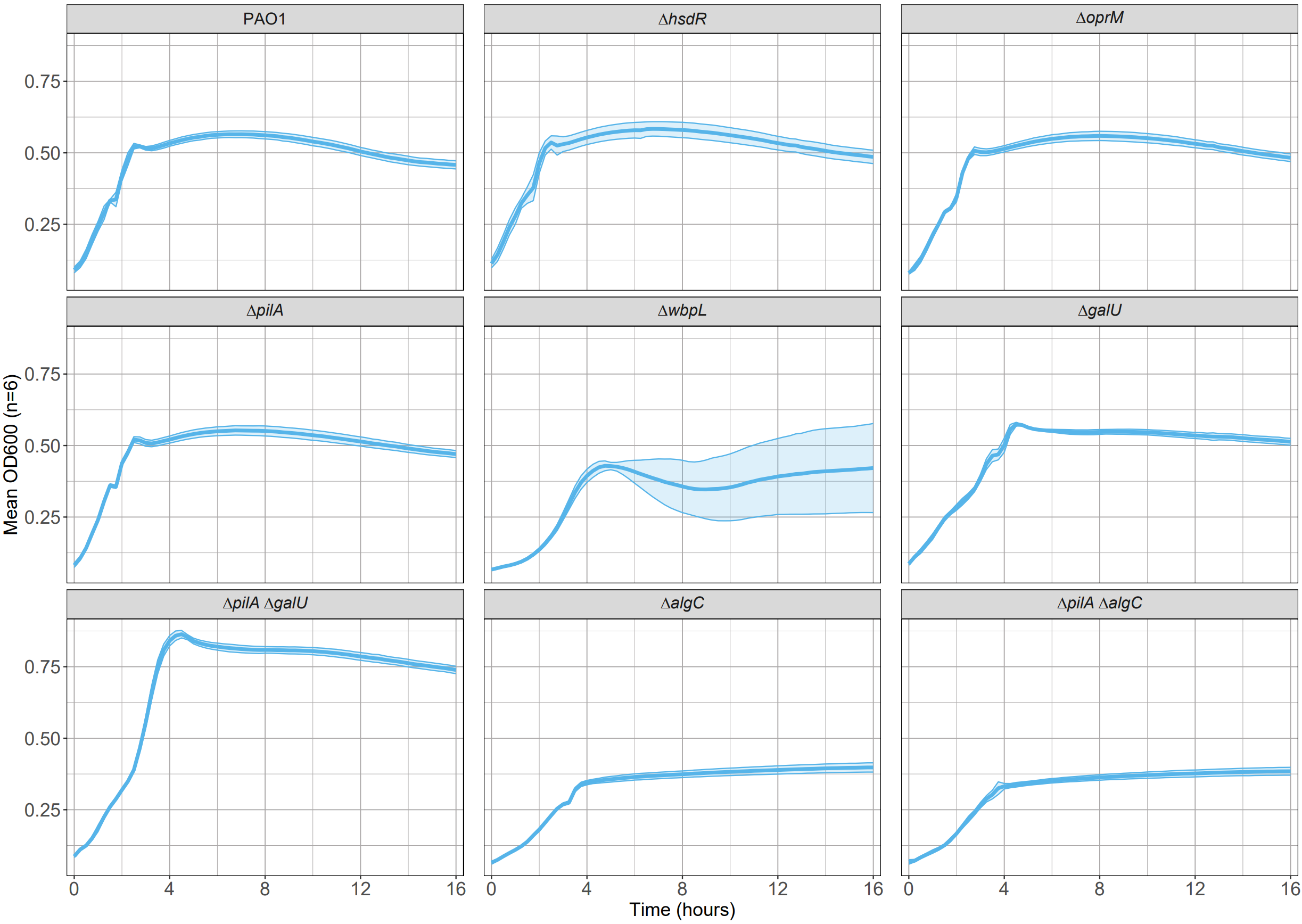

**Supplementary Figure 2:** Mean 16-hour growth curves of the knockout mutants. Ribbons represent 95% confidence interval around the mean (n=6). ∆*hsdR* was used as the parent strain for all subsequent deletions, and ∆*hsdR* is assumed in all the receptor deletion mutants.

|  | **Carrying capacity (K) (+/- 95% CI)** | **Intrinsic growth rate (r) (+/- 95% CI)** | **Doubling time (DT) (hours) (+/- 95% CI)** | **Area under the curve (AUC) (+/- 95% CI)** |
| --- | --- | --- | --- | --- |
| **PAO1** | 0.394 (+/-0.00588) | 2.51 (+/-0.0476) | 0.277 (+/- 0.00517) | 8.97 (+/- 0.129) |
| **∆*hsdR*** | 0.396 (+/-0.0138) | 2.42 (+/- 0.122) | 0.287 (+/- 0.0150) | 9.02 (+/- 0.317) |
| **∆*oprM*** | 0.417 (+/- 0.00937) | 2.14 (+/- 0.0974) | 0.325 (+/- 0.0159) | 9.34 (+/- 0.225) |
| **∆*pilA*** | 0.405 (+/- 0.00775) | 2.59 (+/- 0.122) | 0.268 (+/- 0.0126) | 9.25 (+/- 0.184) |
| **∆*wbpL*** | 0.360 (+/- 0.178) | 2.32 (+/- 0.781) | 0.553 (+/- 0.582) | 7.18 (+/- 2.66) ** |
| **∆*galU*** | 0.426 (+/- 0.0149) | 2.01 (+/- 1.21) | 0.438 (+/- 0.123) | 9.22 (+/- 0.259) |
| **∆*pilA* ∆*galU*** | 0.658 (+/- 0.0146) *** | 2.42 (+/- 0.982) | 0.325 (+/- 0.0733) | 14.25 (+/- 0.304) *** |
| **∆*algC*** | 0.316 (+/- 0.0124) | 1.22 (+/- 0.0703) ** | 0.572 (+/- 0.0339) * | 6.77 (+/- 0.267) ** |
| **∆*pilA* ∆*algC*** | 0.299 (+/- 0.0106) * | 1.32 (+/- 0.0812) ** | 0.527 (+/- 0.0343) | 6.39 (+/- 0.238) *** |

**Supplementary Table 3:** The mean growth parameters of wildtype PAO1 and the panel of knockout mutants determined using Growthcurver^2^. Linear models were used to determine whether each growth parameters differed significantly between knockout mutants and wildtype PAO1 (growth parameter~strain, n=6, ****p*<0.001, ***p*<0.01, **p*<0.05).

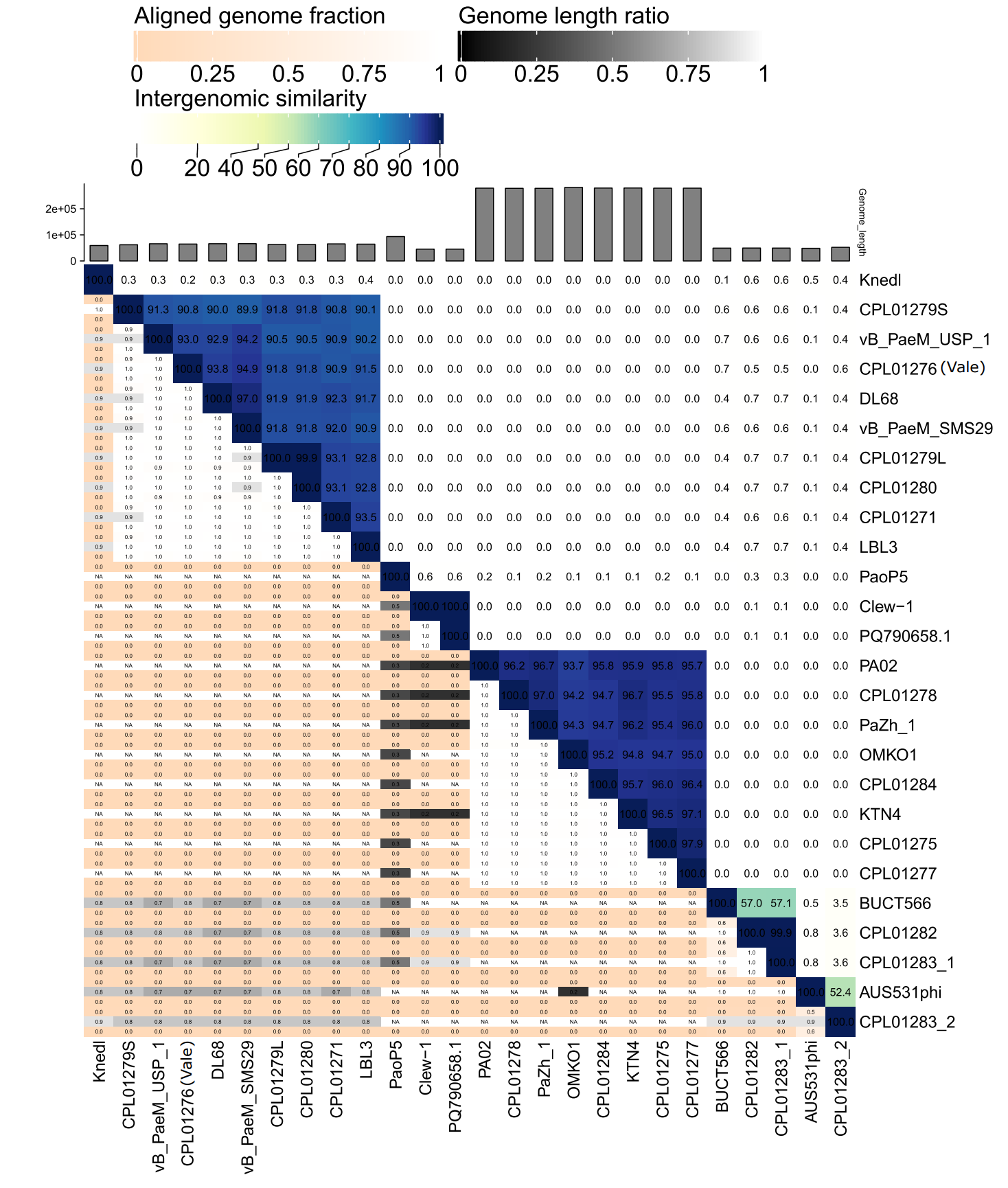

**Supplementary Figure 3:** A VIRIDIC^3^ plot showing the interrelatedness of the phages isolated on ∆*pilA* ∆*galU* in this study, their closest relatives as identified from the NCBI Genbank database using PhageClouds^4^, and other relevant phages discussed in this study. Although discussed in the study, PhiYY was not included in the comparisons as it is a dsRNA virus.

| **Genome** | **Description** | **Accession number** |
| --- | --- | --- |
| Parent strain (∆*hsdR*) | PAO1 with a clean deletion of *hsdR,* lacking the defensive restriction endonuclease. | SAMN54443381 |
| ∆*pilA* ∆galU | PAO1 with clean deletions of *hsdR, pilA* and *galU*, lacking the defensive restriction endonuclease, type IV pili, outer core LPS and O-antigen. | SAMN54443382 |
| Tor_R1 | Tor-resistant mutants generated in a parent strain (∆*hsdR*) background. | SAMN54443383 |
| Tor_R2 |  | SAMN54443384 |
| Tor_R3 |  | SAMN54443385 |
| Vale_R1 | Vale-resistant mutants generated in a ∆*pilA* ∆galU background. | SAMN54443386 |
| Vale_R2 |  | SAMN54443387 |
| Vale_R3 |  | SAMN54443388 |
| CPL01271_R1 | CPL01271-resistant mutants generated in a parent strain (∆*hsdR*) background. | SAMN54549091 |
| CPL01271_R2 |  | SAMN54549092 |

**Supplementary Table 4:** The naïve host and phage-resistant mutant genomes analysed in this study.

| **Affected gene** | **Affected protein** | **Resistant mutant(s) possessing the mutation** | **Type of SNP** | **Nucleotide change** | **Amino acid change** |
| --- | --- | --- | --- | --- | --- |
| *pa2194* (*hcnB*) | Hydrogen cyanide synthase | Tor_R2 | Substitution | 31C>T | G11D |
| *pa5004* (*wapH*) | Putative glycosyltransferase | Vale_R2 | Substitution | 844C>T | P282S |

**Supplementary Table 5:** The SNPs identified in Tor- and Vale-resistant mutants. Nucleotide and amino acid numbers are based on their position in the coding sequence.

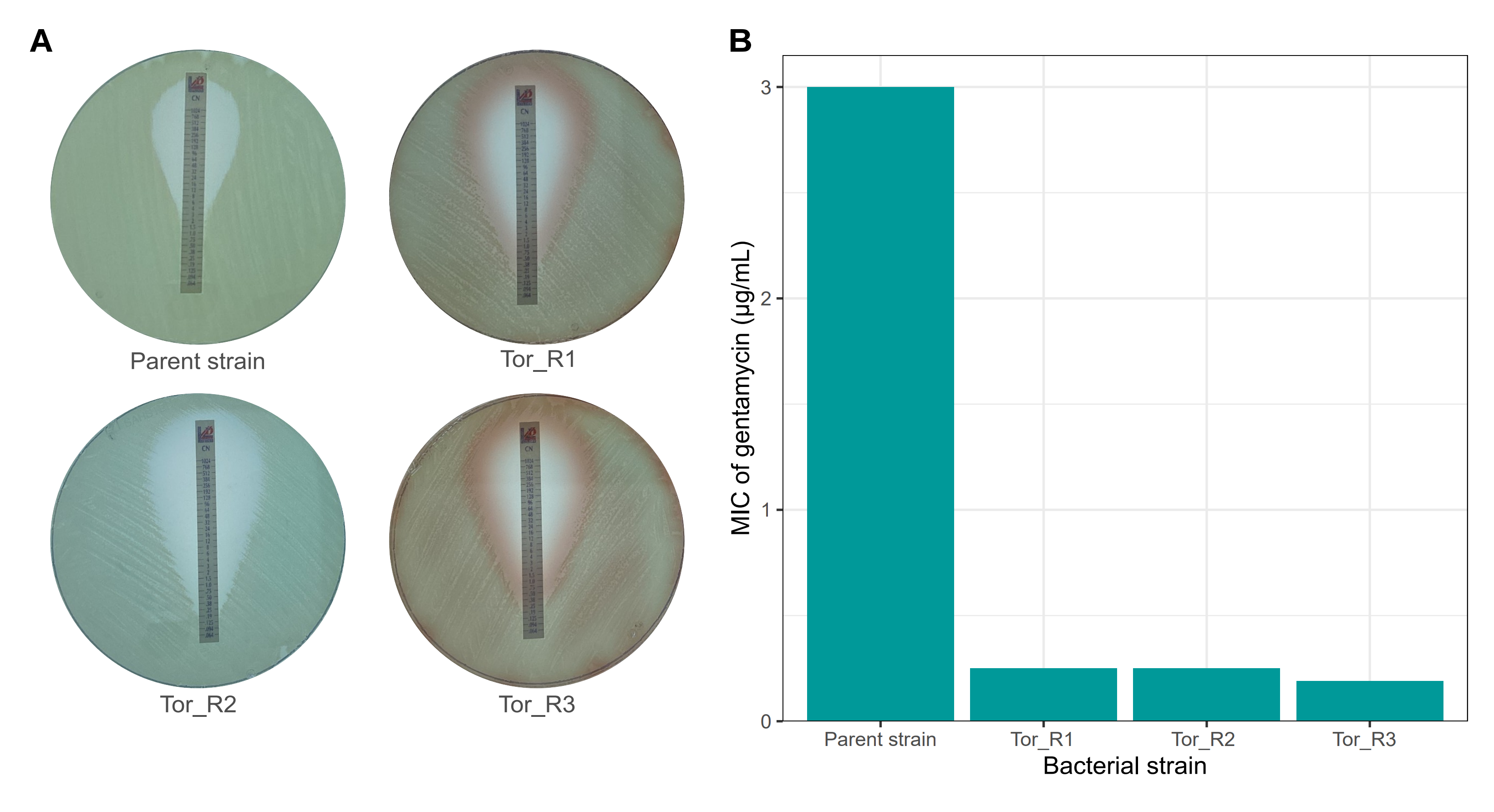

**Supplementary Figure 4: [A]** Liofilm gentamycin test strips applied to lawns of the naïve parent strain and Tor-resistant mutants. **[B]** Phage resistance conferred increased gentamycin suscpetibility in all three resistant mutants, with an MIC of 0.25 µg/mL for Tor_R1 and Tor_R3 and 0.19 µg/mL for Tor_R2, compared 3 µg/mL to for the parent strain*.*

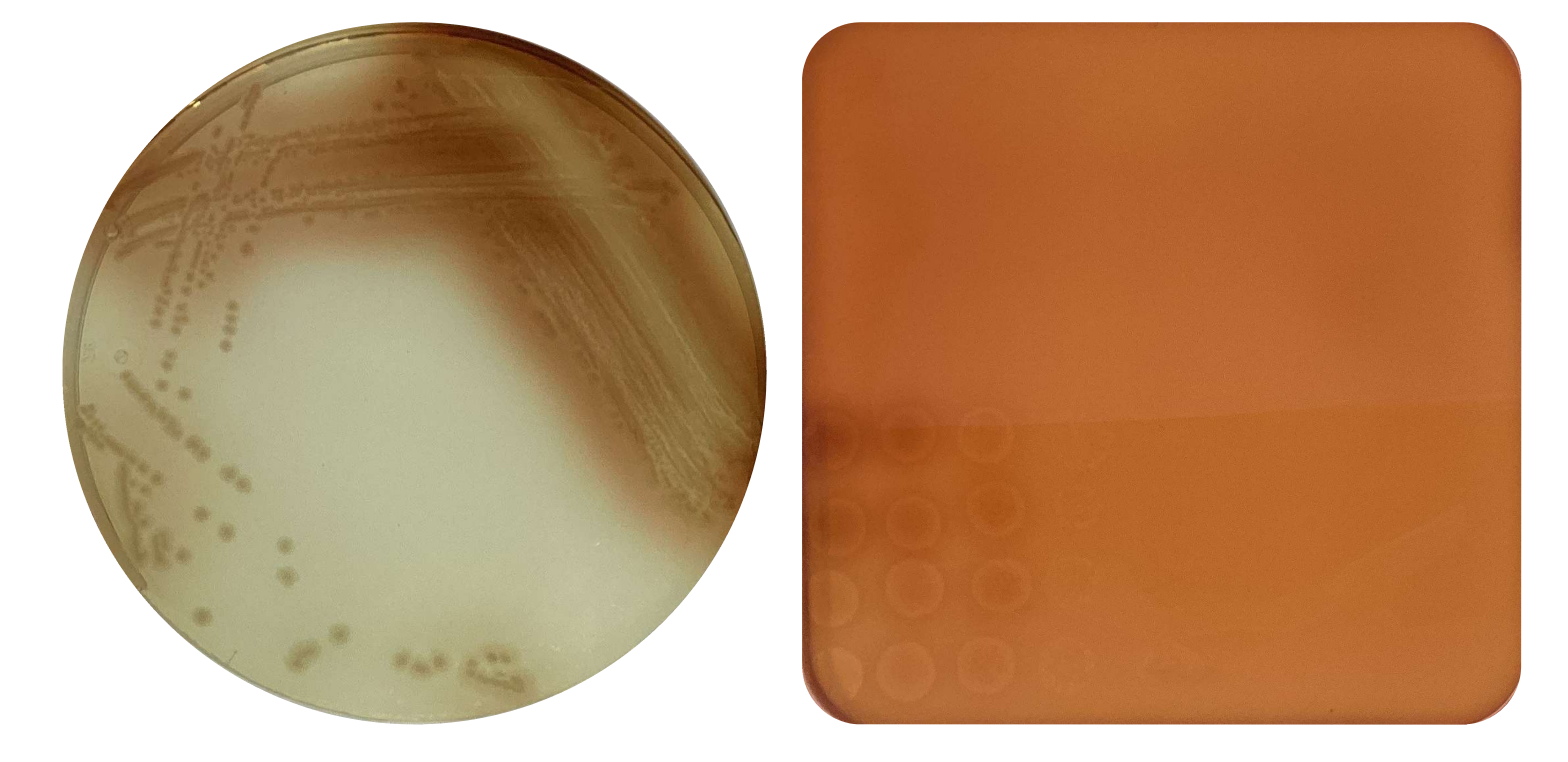

**Supplementary Figure 5:** Tor-resistant parent strain mutants exhibiting a red phenotype associated with the loss of *hmgA* in streak plate (left) and agar overlay plate (right).

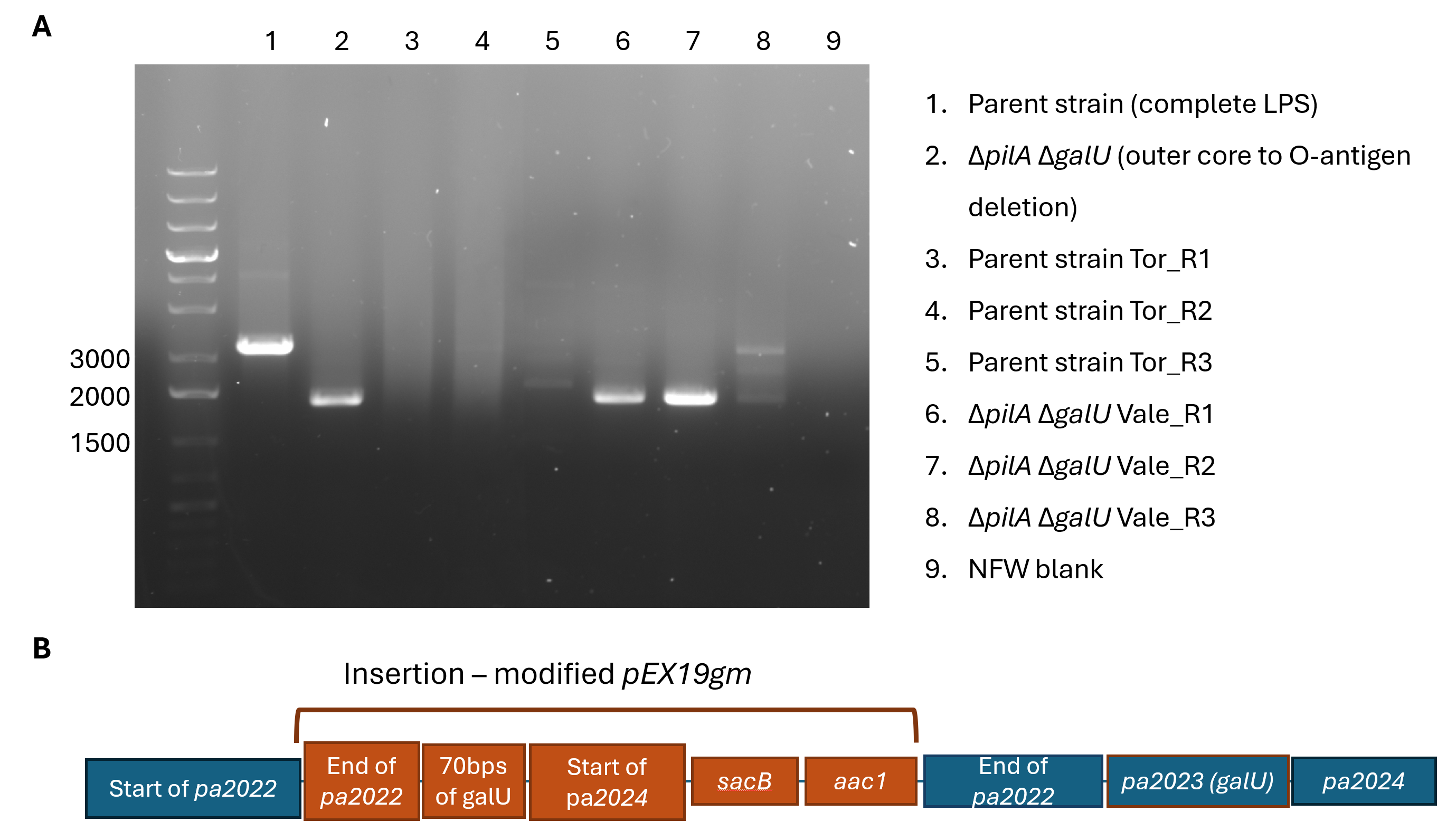

**Supplementary Figure 6: [A]** PCR products of the region surrounding *galU* in phage resistant mutants run on a 1% agarose gel. The parent strain produces a ~2000 bp product*,* while ∆*pilA* ∆*galU,* Vale_R1 and Vale_R2 produce a ~1300 bp product, where *galU* is absent. Tor_R1, Tor_R2 and Tor_R3 produce no band, as the regions upstream and downstream of *galU* where the primers should bind are also absent in the100-200kb deletions. Vale_R3 produces both ~2000 and ~1300 bp products, where wildtype *galU* remains in the genome and the plasmid insert possess the truncated *galU* form in the merodiploid. **[B]** The insertion detected in Vale_R3 was the modified plasmid used for the deletion of *galU,* integrated upstream of the in-tact *galU* gene.
